# Strategies for sampling pseudo-absences for species distribution models in complex mountainous terrain

**DOI:** 10.1101/2022.03.24.485693

**Authors:** Patrice Descombes, Yohann Chauvier, Philipp Brun, Damiano Righetti, Rafael O. Wüest, Dirk N. Karger, Damaris Zurell, Niklaus E. Zimmermann

## Abstract

1. Predictions from species distribution models (SDMs) that rely on presence-only data are strongly influenced by how pseudo-absences are derived. However, which strategies to generate pseudo-absences give rise to faithful SDMs in complex mountainous terrain, and whether species-specific or generic strategies perform better remain open questions.
2. Here, across 500 plant species, we investigated comprehensively how predictions of SDMs at a 93 m spatial resolution are influenced by pseudo-absence strategies, using the complex topography of the Swiss mountains as a model system. We used five generic (random, equal-stratified, proportional-stratified, target, density) and three species-specific (target specific, density specific and geographic specific) approaches to derive pseudo-absence data. We conducted performance tests for each of our eight strategies in combination with (*a*) spatial bias, generated within our occurrence dataset on sites with highest sampling density, to investigate how this common bias problem influences the performance of pseudo-absence sampling strategies, and (*b*) a new approach to reduce model extrapolation in environmental space by including background data from all environmental conditions of the study area. SDMs were evaluated against an independent and well-sampled dataset of true presences and absences.
3. The random, the density (generic), and the geographic specific (species-specific) strategies consistently performed best, even in cases of strong spatial sampling bias in the occurrence data. Including a background of environmentally stratified pseudo-absences improved predictions of species distributions towards environmental extremes, and significantly reduced spatial extrapolations of model predictions in environmental space.
4. Our results indicate that both generic and species-specific pseudo-absence strategies allow estimating robust SDMs and we provide clear recommendations which strategies to choose in complex terrain and when presence data are prone to high sampling bias. In datasets with strong sampling bias, most pseudo-absence strategies produce extrapolation problems and we additionally recommend environmentally stratified pseudo-absences in these cases. Overall, in species rich datasets the use of complex and computationally demanding, species-specific pseudo-absence strategies may not always be justified compared to simpler generic approaches.

## Introduction

Species distribution models (SDMs; Guisan & Zimmermann, 2000; Guisan, Thuiller, & Zimmermann, 2017) are used for many different applications, including predictions of climate change impacts on species distributions (Thuiller et al., 2019) or mapping potential distributions of pests or of threatened species in conservation planning (Araújo et al., 2019; Guisan et al., 2006, 2013; Pearson, 2010). SDMs relate presence and absence records of a species to a set of environmental predictors allowing its ecological niches to be characterized statistically and predicted spatially (Guisan et al., 2017; Guisan & Zimmermann, 2000). Studies have shown that the sampling design for obtaining presence and absence in the landscape may have profound effects on SDM predictions (Chefaoui & Lobo, 2008; Hanberry et al., 2012; Wisz & Guisan, 2009). When sampling is set up before data collection, environmentally stratified survey designs in the field should be preferred over random or gridded designs (Hirzel & Guisan, 2002). This ensures species records that are environmentally balanced across the landscape. However, many studies rely on opportunistically sampled occurrence data from diverse archival or digital sources (e.g., museum datasets, citizen science programs), often gathered without adequate sampling designs, and typically lacking true absence information (Graham et al., 2004). As the majority of SDM algorithms contrast presence data against some form of absence data, many studies have relied on the generation of pseudo-absences. However, different studies have shown that SDM predictions may be strongly affected by the way such pseudo-absences are generated (Chefaoui & Lobo, 2008; Hanberry et al., 2012; Hertzog et al., 2014; Wisz & Guisan, 2009), yet guidance towards adequate pseudo-absence strategies remains limited (Barbet-Massin et al., 2012; Kramer-Schadt et al., 2013; Phillips et al., 2009).

Different pseudo-absence strategies have been proposed to provide alternatives to true absences, ranging from fully random to environmentally stratified approaches, and often including complex spatial manipulations (e.g. Chefaoui & Lobo, 2008; Stokland, Halvorsen, & Støa, 2011; Barbet-Massin, Jiguet, Albert, & Thuiller, 2012; Hanberry et al., 2012; Kramer-Schadt et al., 2013; Senay, Worner, & Ikeda, 2013; Hertzog, Besnard, & Jay-Robert, 2014; Iturbide et al., 2015; Chapman, Pescott, Roy, & Tanner, 2019). Pseudo-absences selected randomly or from environments similar to the occurrence records have already demonstrated gains in SDM performances (Barbet-Massin et al., 2012; Kramer-Schadt et al., 2013) and the most constrained predictive distribution maps (Chefaoui & Lobo, 2008; Hanberry et al., 2012). In contrast, pseudo-absences selected far from species occurrences, often tend to over predict species distributions despite showing good performance indices overall (Chefaoui & Lobo, 2008; Hanberry et al., 2012). Such a pseudo-absence strategy, often referred to as the “unsuitable background concept”, increases the distinctiveness with regards to occurrences (Chapman et al., 2019; Thuiller et al., 2004). However, manipulating pseudo-absence selection by excluding sampling in areas of observed presences has been criticized, because this involves *a priori* assumptions about habitat suitability before actually quantifying it (Stokland et al., 2011). Increasing the spatial or environmental distance between occurrences and pseudo-absences might also simplify the models to a few main environmental drivers (VanDerWal et al., 2009). In addition to generic, also species-specific pseudo-absence generation approaches have been proposed. However, neither comparative accuracies nor computational costs of applying species-specific vs. generic pseudo-absences strategies have been addressed so far. In summary, there is still no clear consensus on which pseudo-absence sampling strategies should be implemented in SDMs, and especially for species rich datasets and complex terrain.

Overall, SDMs based on spatially biased occurrence datasets show improved performance if a similar spatial bias is introduced for sampling pseudo-absences as is found in presences (Elith & Leathwick, 2007; Hertzog et al., 2014; Kramer-Schadt et al., 2013; Phillips et al., 2009). For instance, Kramer-Schadt et al. (2013) and Hertzog et al. (2014) used the density of occurrences to randomly select pseudo-absences across the landscape, resulting in a higher number of pseudo-absences in regions with a higher density of species observations. Similarly, the target group approach (Phillips et al., 2009), uses all occurrences of a pre-defined species group (the target group) collected with the same sampling design as the species of interest, was shown to improve the quality of model predictions (Elith & Leathwick, 2007; Phillips et al., 2009). However, the definition of a target group involves an arbitrary decision on membership to the target group and the ecology of the species. For example, the group selection might focus on a clade, a family, or on some functional characteristics. Kramer-Schadt et al. (2013) also recommended that spatial thinning of occurrences should be preferred over bias compensation with pseudo-absences, if a sufficiently high number of occurrences is available. While pseudo-absence strategies have been proposed for highly biased datasets, and for coarse spatial resolution studies, an overview of the best sampling strategies in complex terrain and at high spatial resolution (i.e. < 1 km) is missing.

Assessing the accuracy of spatial extrapolations of species predictions into environmental conditions not covered by presence and absence data (Mesgaran et al., 2014) has received limited attention so far when considering different pseudo-absence sampling strategies. For instance, *Androsace helvetica* L., a cushion plant species growing at very high elevations in the European Alps (i.e., 1800-3200 m a.s.l.), might be predicted above its known upper elevation range limit if no pseudo-absences are selected at higher elevations than where it is known to occur. Missing pseudo-absences in these regions could in fact result in truncated or incomplete model response curves, leading to unrealistic spatial predictions in areas where the species neither grows nor reproduces (Thuiller et al., 2004; Zurell et al., 2012). Moreover, the currently favoured pseudo-absence strategies – i.e., target-group and sampling density-based approaches – may not avoid such extrapolation issues into marginal environments that are not well covered by the presence observations. These strategies indeed tend to frequently select low numbers of pseudo-absences in environments with low density of presenses, because of a generally lower sampling effort in these regions. If SDM-based projections are used to define suitable areas for conservation purposes or species re-introductions, model overpredictions might then introduce incorrect guidance to field biologists, environmental practitioners and stakeholders. To avoid this problem, considering environmental stratification for generating pseudo-absences has been proposed (Thuiller et al., 2004). Yet, we lack a clear overview of how such stratification affects projected species distributions when combined with different pseudo-absence sampling strategies.

Here, we investigated how predicted species occurrences from SDMs (ensemble averaging of five algorithms) are influenced by pseudo-absence sampling strategies at high spatial resolution (i.e. 93 m) for 500 plant species across the complex topography of Switzerland. Eight pseudo-absence strategies were implemented and combined with additional sets of environmentally stratified background points, so that pseudo-absence points cover the entire environmental space of our study area for model calibrations (Fig. 1). Spatial bias in the occurrence dataset was additionally manipulated to investigate how the spatial presence sampling bias influences the performance of pseudo-absence sampling strategies. Predictions from a single model, and ensemble means of all models were evaluated against an independent dataset of true presences and absences that samples the whole study area and environmental gradients well. Finally, the degree of extrapolation to novel environments was evaluated, the performance of the eight different pseudo-absence sampling strategies compared, and recommendations are provided. Our aim here is to identify which pseudo-absence strategy is best suited to model large sets of species in complex terrain, and to list a set of recommendations that may be applied across a wider range of organisms groups and study areas.

**Figure 1.**
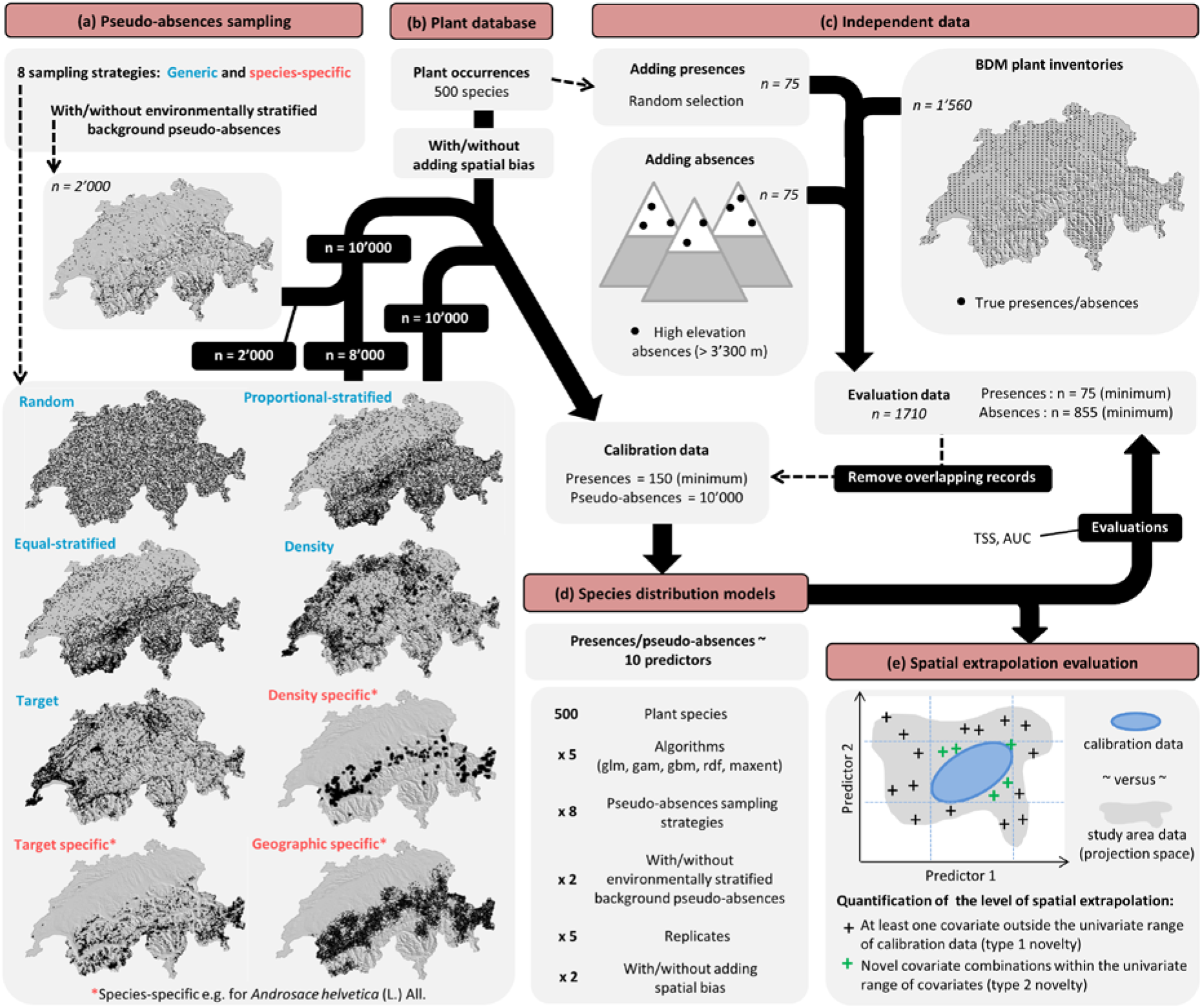
Overview of the method, databases and main analyses performed in the present study. Example maps in the lower left figure are illustrated for *Androsace helvetica* (L.) All.

## Methods

### Plant databases

We used plant records from the National Data and Information Center on the Swiss Flora (Info Flora: www.infoflora.ch, data extracted in April 2019), containing 6.7 million records of 4’238 plant taxa. We retained geographically valid occurrences with a coordinate precision < 100 m (5.1 million records of 4’057 plant taxa), with most of the records (95%) sampled after 1985. We then selected 500 plant species (incl. aggregates or subspecies) that are native to Switzerland, span different ecological groups according to Flora Indicativa (Landolt et al., 2010), and retained at least 250 occurrences after a 300 m spatial thinning. Species with ≥ 250 observations were chosen to avoid sample size effects (Hernandez et al., 2006; Wisz et al., 2008; Zimmermann et al., 2007), and to achieve a presence-to-predictor ratio ≥ 10 per species (Guisan et al., 2017; Harrell et al., 1996). Spatial thinning was performed to avoid pseudo-replication of records, and a 300 m threshold was defined because it significantly reduced spatial autocorrelation in residuals of both generalized linear models (glm; McCullagh, 1983) and generalized additive models (gam; Hastie & Tibshirani, 1987; Guisan, Edwards, & Hastie, 2002) in preliminary analyses (See Note S1, Fig. S1 and S2 in supplementary material for details).

Plant inventories of the Swiss biodiversity monitoring survey (BDM indicator Z9 ‘species diversity in habitats’, www.biodiversitymonitoring.ch; Fig. 1) were used as an independent validation dataset. The dataset spanned the years 2000 to 2018 and consisted of 1’560 circular exhaustive plant inventories of 10 m^2^ evenly spread over Switzerland. We considered all recorded plants across all replicated inventories for each plot location, and species not recorded on plot locations were considered absent.

For generating sampling bias in the occurrence dataset, we used a polygon shapefile of the national floristic sampling campaign defined by Welten & Sutter (1982) and consisting of 593 polygons (mean area = 69.6 km^2^; area range = 0.1-231 km^2^) distributed across Switzerland. These polygons were delimited by experts and were based on political and watershed boundaries as well as the treeline (Fig. S3).

### Environmental predictors

We used a digital elevation model (DEM) at 93 m resolution (Robinson et al., 2014) as base map to derive 13 different topographic, landuse, geological and climate predictors. Ten of these predictors were used for modelling, namely: topographic position index (TPI), topographic roughness index (TRI), topographic wetness index (TWI), mean annual temperature (Tave year), annual precipitation sum (Prec year), annual sum of solar radiation (Srad year), average normalized difference vegetation index (NDVI mean), inner forest density (Forest height Q25), soil humidity (EIV-F) and soil pH (EIV-R). Three additional predictors were used to construct pseudo-absence sampling strategies: elevation, slope and aspect.

Elevation was taken from the 93 m DEM (Robinson et al., 2014). Slope, aspect, TPI and TRI were calculated with the *terrain* function from the *raster* package (Hijmans, 2019) in R 3.5.1 (R Core Team, 2018), and TWI was calculated with SAGA GIS (Conrad et al., 2015), by using the DEM. Tave year, Prec year and Srad year (direct and diffuse potential solar radiation) layers were retrieved from Zimmermann & Kienast (1999) at 25 m spatial resolution and aggregated to 93 m using the average and reprojected to our projection system using bilinear interpolation. NDVI mean was calculated by using Landsat data (https://espa.cr.usgs.gov/) collected for the years 2007-2015 from 1 July to 15 September at a 30 m resolution and aggregated to 93 m using bilinear interpolation. Inner forest density was estimated by analysing cantonal and Swisstopo (www.swisstopo.admin.ch) LiDAR data acquired between 2000 and 2014 with a density of 0.5–35 points/m^2^ (Artuso et al., 2003; Wüest et al., 2020). We removed the information on buildings by using the Topographic Landscape Model (TLM from Swisstopo) and normalised the returns with the digital terrain model siwssAlti3D from Swisstopo. We calculated the 25^th^ height-percentile of returns above 40 cm as a measure of inner forest density (Wüest et al., 2020) at a spatial resolution of 25 m, and aggregated it at 93 m using bilinear interpolation. EIV-F and EIV-R were retrieved from Descombes et al. (2020). These are proxies of soil wetness and pH, respectively, derived by spatial modelling of plant ecological indicator values (EIVs; Landolt et al., 2010) and have been shown to be important predictors of plant distribution in SDMs (Descombes et al., 2020).

### Pseudo-absence sampling strategies

To investigate how pseudo-absence sampling strategies affect predictions of species distributions, we selected 10’000 pseudo-absences following five generic (i-v) and three species-specific (vi-viii) sampling strategies (Fig. 1).

i. *Random*. Pseudo-absences are selected spatially at random across the study area in Switzerland. This allows the sampling to be randomly distributed across the landscape.
ii. *Equal-stratified*. We reclassified slope, aspect and elevation layers separately into 9 value bins with the *ecospat*.*rcls*.*grd* function from the *ecospat* package (Broennimann et al., 2020; Di Cola et al., 2017), and combined them into unique combinations of these three factors along their full environmental range (n = 84 strata). We then randomly selected, whenever possible, an equal number of pseudo-absences per stratum with the *ecospat*.*recstrat_regl* function from the *ecospat* package (Broennimann et al., 2020; Di Cola et al., 2017). This sampling strategy is topographically stratified and refers to the equal-stratified sampling design described in Hirzel & Guisan (2002). It allows the topographic space to be equally sampled across the study area and guarantees that all strata are represented in topographically complex terrain irrespective of how abundant the strata are.
iii. *Proportional-stratified*. Pseudo-absences were randomly selected proportionally to the logarithm of the area of each topographic stratum with the *ecospat*.*recstrat_prop* function from the *ecospat* package (Broennimann et al., 2020; Di Cola et al., 2017). This strategy refers to the proportional-stratified sampling design described in Hirzel & Guisan (2002). It allows the topographic space to be sampled proportional to the area of the strata across the study area, while guaranteeing that rare strata are still represented.
iv. *Density*. We calculated at a 5 km resolution the density of all plant occurrences (5.1 million records of 4’057 plant taxa) across Switzerland, by dividing the total number of occurrences per 5 km cell by the total number of occurrences in Switzerland. We then used this density layer as a vector of probability weights to sample pseudo-absences. This results in a higher likelihood of sampling pseudo-absences in regions of higher overall sampling effort. This sampling strategy mimics the sampling bias of presences, resulting in sampling of pseudo-absences more often in the lowland and close to cities (where citizen scientists recorded most plant occurrences). It also avoids placing pseudo-absences in regions where no presences were sampled.
v. *Target*. We used all plant occurrences as potential pseudo-absences following Phillips et al. (2009). This sampling strategy, often called “target group” approach, is very similar to the density approach and re-samples pseudo-absences from the pool of observations of presences irrespective of species identity.
vi. *Density specific*. We calculated at a 5 km resolution the occurrence density of the species of interest across Switzerland, by dividing its number of occurrences per 5 km cell by its total number of occurrences in the country. We then used this density layer as a vector of probability weights, to sample more pseudo-absences in regions with higher sampling efforts for the species of interest. This strategy allows regional sampling bias to be considered by selecting more pseudo-absences from regions where the species of interest is sampled more frequently.
vii. *Target specific*. Pseudo-absences were selected within the basic elevation range (± 10% of the observed elevation range) of the species of interest. For this, elevation values of all records were extracted and the minimum and maximum elevation of each plant species (n = 4’057) was calculated. We then retained the presences of all species occurring within the so-defined elevation range of the species of interest, i.e., ± 10% of the elevation range, and used their occurrences as potential pseudo-absences. This sampling strategy selects only pseudo-absences from cells that were visited at least once for any other species in close elevational proximity to the species of interest. It thus selects potential pseudo-absences from species that inhabit more similar elevation ranges compared to the generic target approach.
viii. *Geographic specific*. Pseudo-absences were here selected in regions where the species of interest had been observed. For this, an interpolation-based model was constructed to predict the probability of being close to an occurrence of the species of interest across Switzerland. Inverse distance weighting interpolation from the *geoIDW* function in the *dismo* R package (Hijmans et al., 2017) were used on equal numbers of observed presences and (spatially) randomly generated absences of the species of interest to build the interpolation surface. This surface was then used to randomly sample pseudo-absences proportional to the probability of presence. This approach is very similar to the density specific sampling strategy, but is more computationally demanding and the sampling intensity is not higher in areas that are more frequently sampled for the species.

For each design, we sampled 10’000 pseudo-absences at a 93 m resolution across Switzerland and each design was repeatedly sampled five times (5 replicates). To reduce spatial autocorrelation in model residuals, pseudo-absences were selected such that a minimum distance of 300 m among sampled locations was guaranteed. We allowed pseudo-absences at or close to presence locations following the recommendations of Stokland et al. (2011). Indeed, artificially removing pseudo-absences in the proximity of occurrences represents a strong arbitrary assumption, and more generally, absence observations of a species in regions of potentially suitable areas are not contradictory, but a natural phenomenon.

### Introducing sampling bias in species occurrences

To investigate how sampling bias in species occurrences might affect the choice of the best pseudo-absence strategy, we sub-sampled the occurrence data sets to include only presences from the 10% most sampled regions. Regions were delineated using spatial information from the national mapping program of Welten & Sutter (1982). For every species, we randomly selected 150 occurrences from the (spatially thinned) occurrence data sets that were located within the 10% most sampled Welten & Sutter polygons (see Fig. S3 for an example). If 150 occurrences were not available within the 10% most sampled Welten & Sutter polygon areas, we allowed higher percentages (e.g., 15% most sampled polygons) until at least 150 occurrences per species were sub-sampled. By this, we constructed occurrence datasets that only sample species observations in the most sampled regions relative to the species of interest. We then repeated the species-specific pseudo-absence data selection for the newly generated set of biased occurrences, while the pseudo-absence sets generated with a generic approach did not require updates.

### Adding environmentally stratified background data to pseudo-absences

In order to assess the importance of covering the whole environmental space of the study area with the pseudo-absence set, we sampled an additional 2000 points in an environmentally stratified manner based on the predictors used in the SDM models (see below). To this end, we reclassified each of the 10 environmental predictors used for modelling into 5 equally spaced bins with the *ecospat*.*rcls*.*grd* function from the *ecospat* R package (Broennimann et al., 2020; Di Cola et al., 2017), and combined them into composite environmental strata that represent unique combinations of the individual ten classified predictors along their full environmental range (n = 2088 strata). From the output layer, 2’000 pseudo-absences were then randomly sampled (1 per stratum) using the *ecospat*.*recstrat_regl* function from the *ecospat* R package (Broennimann et al., 2020; Di Cola et al., 2017). Pseudo-absences were also sampled with a minimum distance of 300 m between sampled points, as we did for occurrences. The sampling was repeated five times (5 replicates). This approach allows the entire environmental range to be covered by the pseudo-absence data, ensuring that modelled probabilities do not extrapolate into parts of the environmental space not covered by the calibration dataset. This background of environmentally stratified pseudo-absences (hereafter called “Background strata”) replaced (or not) for each pseudo-absences sampling strategy 2’000 random pseudo-absences generated by the eight different methods (Fig. 1), since these established methods do not explicitly avoid such extrapolations.

### Species distribution models

We used species distribution models (SDMs; Guisan & Zimmermann, 2000; Guisan et al., 2017) to relate each set of occurrences and pseudo-absences to 10 environmental predictors representing current topographic, landuse, geological and climate conditions. Our 10 selected environmental predictors were chosen due to their known general influence on plant species distributions (Descombes et al., 2020). They showed pairwise Pearson correlations |r| < 0.6, being well within the tolerable range suggested by Dormann et al. (2013). We selected no more than 10 predictors, since model performance has been demonstrated to peak around 10-11 predictors (Brun et al., 2020).

We modelled each species following an ensemble approach of different statistical techniques (Araújo & New, 2007) using five different model algorithms: generalized linear models (glm; McCullagh, 1983), generalized additive models (gam; Hastie & Tibshirani, 1987; Guisan et al., 2002), gradient boosting machines (gbm; Ridgeway, 1999; Friedman, 2001), Random Forests (rdf; Breiman, 2001) and maximum entropy models (MaxEnt; Phillips, Anderson, & Schapire, 2006; Elith et al., 2011). Ensemble models tend to show better performances than single algorithm models (Liu et al., 2019). We gave equal weights to occurrences and pseudo-absences in model calibrations and models were repeated five times with the 5 replicated sets of pseudo-absences (Barbet-Massin et al., 2012; Liu et al., 2019; Wisz & Guisan, 2009).

Models were evaluated against the independent BDM dataset. Since several species had less than 50 occurrences in the BDM dataset, we randomly shifted 75 occurrences per species from the training to the evaluation dataset to ensure a robust evaluation of the models. In addition, because few BDM plant inventories were performed at very high elevation (for technical and security reasons), we also added 75 absences above 3’300 m, an elevation above which hardly any species is known to grow and reproduce in the Swiss Alps (Körner, 2003). All species in the database of the National Data and Information Center on the Swiss Flora have their 97.5^th^ percentile of the elevation distribution below this limit. Models were calibrated without the grid cells containing occurrences or absences of the independent dataset, ensuring that the evaluation dataset is fully independent from the calibration dataset. We evaluated the ability of the model to discriminate occurrences from absences with the True Skill Statistics (TSS; Allouche, Tsoar, & Kadmon, 2006) and the area under the ROC-plot curve (AUC; Fielding & Bell, 1997). We converted model probabilities to simulated presences and absences by using the threshold maximising the TSS and AUC and by using functions implemented in the *PresenceAbsence* R package (Freeman & Moisen, 2008). Finally, evaluation metrics were averaged across replicates for every algorithm and species, and an ensemble evaluation was performed by assessing average predictions calculated across replicates and algorithms for every species. Model performances were compared by pairwise comparisons with paired Wilcox tests and displayed following a letter-based representation of significant difference and a Bonferroni correction was applied for multiple tests.

### Evaluating extrapolation into novel covariate space

Because differences in model extrapolation to novel covariate combinations in environmental space may arise across different pseudo-absence strategies, we quantified the level of model extrapolation for each calibration dataset (i.e., environmental coverage of the calibration data versus projected space). We quantified the level of model extrapolation to new environmental space, for geographic regions that have at least one covariate outside the univariate range of calibration data (type 1 novelty), and regions that represent novel covariate combinations within the univariate range of covariates (type 2 novelty) following (Mesgaran et al., 2014). We calculated the percentage of cells concerned by type 1 novelty (NT1; values < 0) and type 2 novelty (NT2; values > 1), and determined the most related influential covariates. Finally, for each species, we retained cells presenting NT1 and NT2, and summed them across species for both NT1 and NT2 separately.

## Results

### Model evaluations

Random forest showed on average the best predictive performance compared to other algorithms although differences were small overall (average difference: TSS = +0.020, AUC = +0.010; Table S1 and S2). Ensemble models showed better average evaluation performance than averaged single models (average difference: TSS = +0.013, AUC = +0.008; Table S1 and S2). For this reason, all subsequent analyses were interpreted on the basis of the ensemble model. Overall, ensemble models showed good predictive performances under all pseudo-absence sampling strategies (TSS >0.4, AUC >0.7), except for a few species and when sampling bias was added to the species occurrence datasets (Fig. 2 and S4).

**Figure 2.**
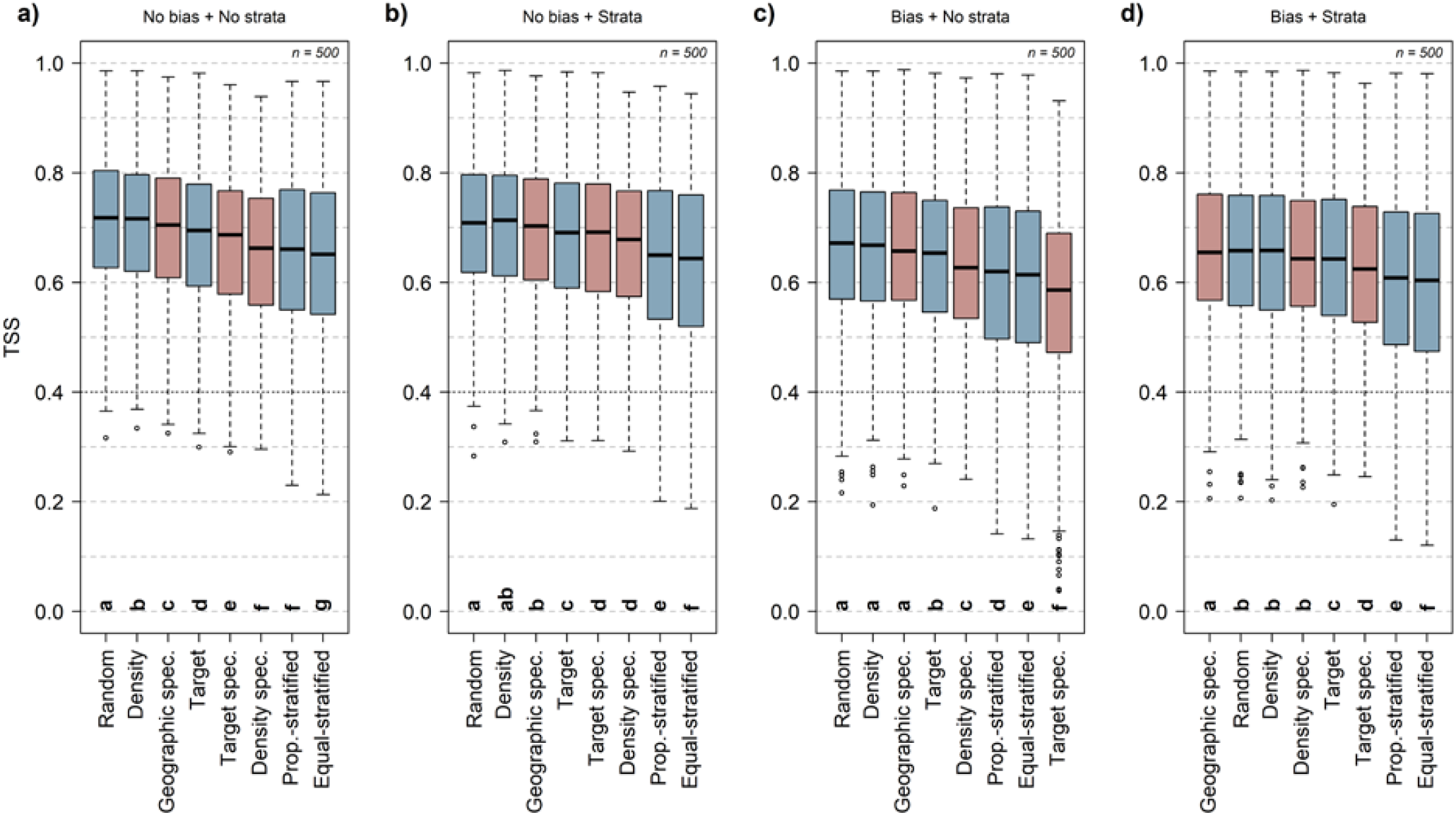
SDM performance (TSS metric) of ensemble models assessed across 500 species (average of replicates) under eight pseudo-absences sampling strategies (blue = generic; red = species-specific). Models were run with the original thinned set of occurrences (a, b; “No bias”) or with a spatially biased subset of occurrences (c, d; “Bias”). Pseudo-absences were sampled following the eight pseudo-absences sampling strategies and included (b, d; “Strata”) or did not include (a, c; “No strata”) an environmentally stratified background of pseudo-absences in the calibration dataset. The pseudo-absence sampling strategies are ranked according to their average performances (high to low from left to right). The dotted black line represents the thresholds above which models are considered to have reliable performances (TSS > 0.4). All pairwise comparisons were run with paired Wilcox tests with Bonferonni correction for multiple tests and displayed following a letter-based representation of significant difference (p-value > .05).

### Selecting the best pseudo-absence strategy

The random, the generic density and the geographic specific pseudo-absence strategies ranked first, second and third by predictive performance of the ensemble models, respectively, when no sampling bias in species occurrences, or background of environmentally stratified pseudo-absences, was added (Fig. 2a, S4, Table S1 and S2). Adding sampling bias to species occurrences decreased overall model performance of ensemble models (average difference: TSS = -0.048, AUC = -0.025; Fig. 3a, S5, Table S1 and S2). Yet, adding bias did not change the rank order among the best three pseudo-absence strategies (Fig. 2c and S4). The target specific pseudo-absence approach showed the highest drop in model performance when a sampling bias was added to the species occurrence data (average difference: TSS = -0.104, AUC = -0.060; Fig. 3a, S5, Table S1 and S2).

**Figure 3.**
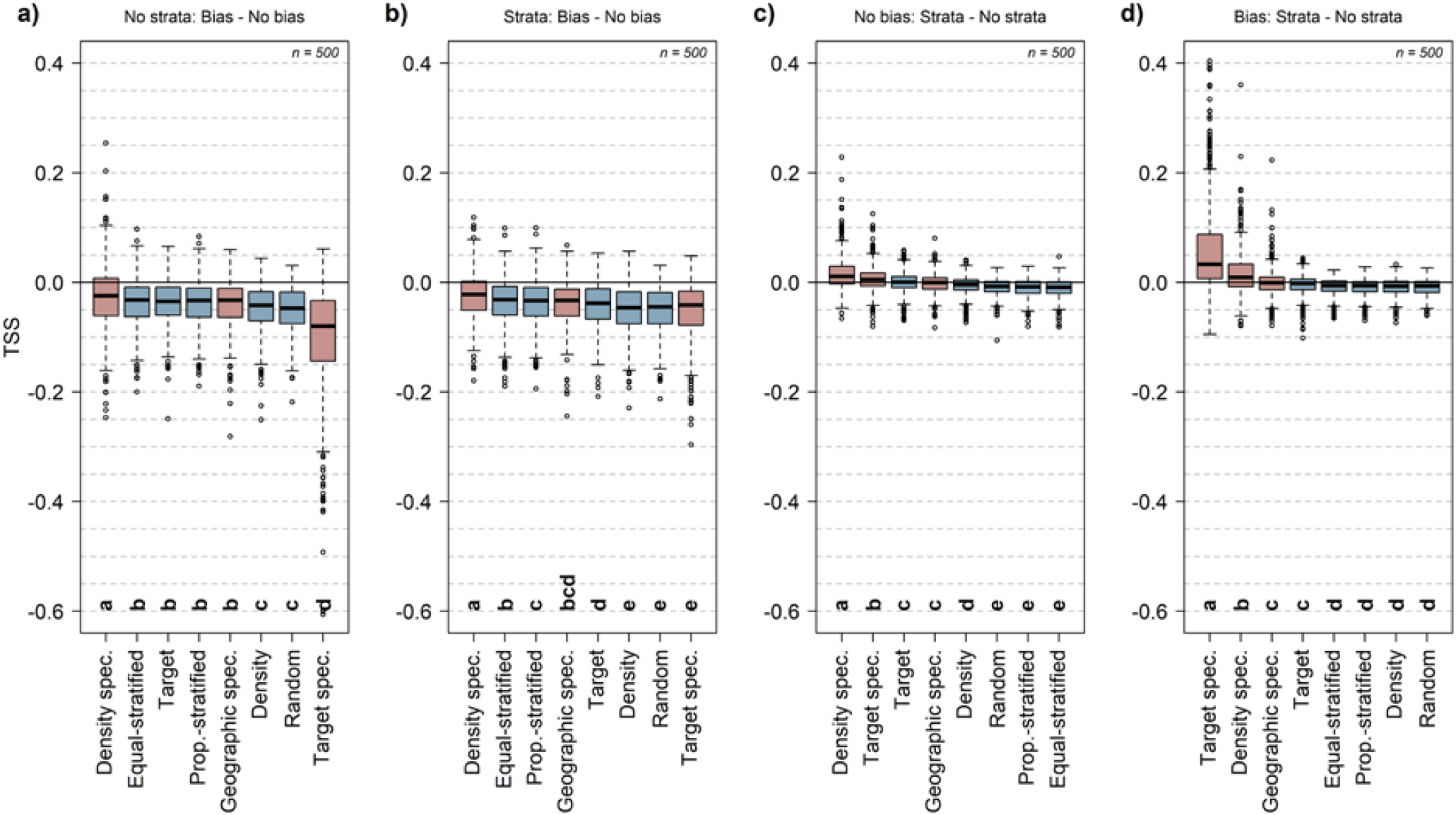
Pairwise differences in SDM performance (TSS metric) of ensemble models assessed across 500 species (average of replicates) under eight pseudo-absences sampling strategies (blue = generic; red = species-specific) in response to manipulations of pseudo-absence data. *Panel a* and *b* represent the effect of implementing spatial bias in the set of occurrences on the model performance (“Bias – No bias”) when the set of pseudo-absences did not include (a; “No strata”) or included (b; “Strata”) an environmentally stratified background of pseudo-absences in the calibration dataset. *Panel c* and *d* represent the effect of including an environmentally stratified background of pseudo-absences in the pseudo-absences sets (“Strata – No strata”) when the models were calibrated with the original thinned set of occurrences (c; “No bias”) or with a spatially biased subset of occurrences (d; “Bias”). The pseudo-absence sampling strategies are ranked according to their average difference in performances (high to low from left to right). All pairwise comparisons were run with paired Wilcox tests with Bonferonni correction for multiple tests and displayed following a letter-based representation of significant difference (p-value > .05).

### Adding a background of environmentally stratified pseudo-absences

The performance of ensemble models based on the target specific and density specific pseudo-absence strategies generally improved when an environmentally stratified background of pseudo-absences was added to the calibration data; both without (average difference: density specific, TSS = +0.016, AUC = +0.008; target specific, TSS = +0.007, AUC = +0.004; Fig. 3c, S5, Table S1 and S2) and with added spatial bias in the occurrence dataset (density specific, TSS = +0.017, AUC = +0.011; target specific, TSS = +0.059, AUC = +0.039; Fig. 3d, S5, Table S1 and S2). In particular, the target specific pseudo-absence strategy, combined with spatially biased occurrences, showed high improvements in model performance when a background of environmentally stratified pseudo-absences was added (average difference: target specific, TSS = +0.059, AUC = +0.039; Fig. 3d, S5, Table S1 and S2). The performance of the other pseudo-absence strategies was insensitive to these manipulations or decreased slightly (average difference: TSS = -0.011 to 0.000, AUC = -0.005 to 0.000; Fig. 3c, 3d, S5, Table S1 and S2). The density specific, target specific and geographic specific pseudo-absence strategies showed distinct improvements in ensemble model performance for a few species (species with average difference in TSS > +0.05 or AUC > +0.05; Fig. 3c, 3d, and S5).

Overall, the ranking of the best pseudo-absence strategies was the same when the unbiased, original calibration data was used compared to using an environmentally stratified background of pseudo-absences in the calibration data (rank 1 to 3: random, density and geographic specific, respectively; Fig. 2b, S4, Table S1 and S2). However, and despite marginal differences, the geographic specific sampling strategy performed better than the random and the generic density strategies when occurrences were spatially biased (Fig. 2d, S4, Table S1 and S2).

### Spatial extrapolation into novel covariate space

Model extrapolation into novel covariate space varied between the different pseudo-absence sampling strategies (Fig. 4). When no background of environmentally stratified pseudo-absences or bias in occurrences were added, type 1 novelty (NT1) model extrapolation in space was low for random (median value: 0.14%), proportional-stratified (0.23%), equal-stratified (0.30%) and density (0.60%) pseudo-absence sampling strategies, but presented high median values or high variability for target (1.87%), density specific (2.20%), target specific (8.89 %) and geographic specific (0.52%) pseudo-absence strategies (Fig. 4a and S6). Type 2 novelty (NT2) model extrapolation in space was low for all pseudo-absence strategies (median values: < 0.29%), but presented high variability and peaked for some species for density specific (max value: 27.84%), target specific (15.30%) and geographic specific (12.53%) pseudo-absence strategies (Fig. 4b and S7). Overall, NT1 and NT2 mostly occurred in high elevation sites presenting temperatures below the range of the calibration data (i.e., >2800 m) and wetlands such as lakes which are poorly represented in the calibration data (Figs. S6, S7, S8 and S9).

**Figure 4.**
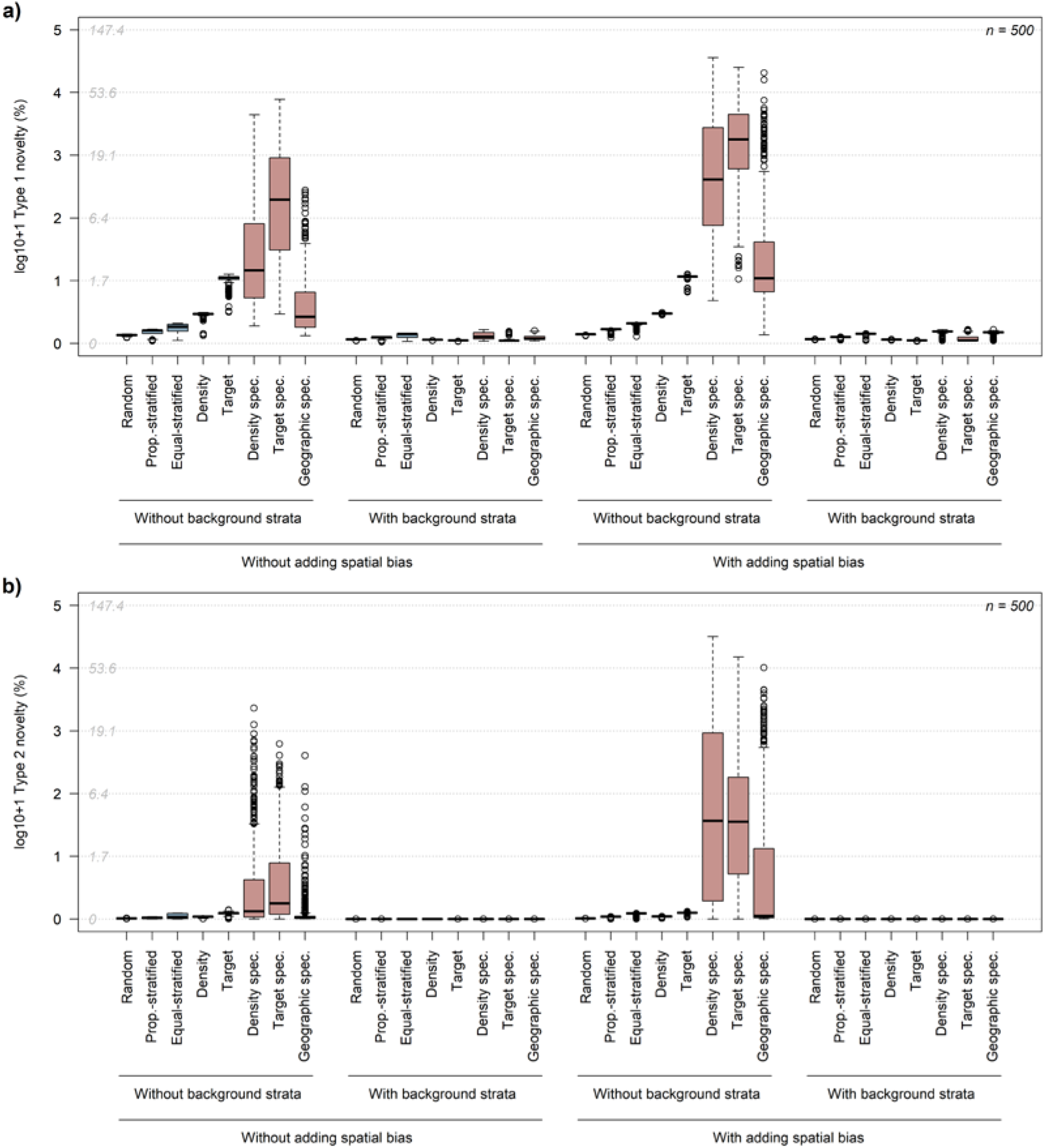
Quantification of the level of spatial extrapolation from the calibration data (presences and pseudo-absences) in the projection space (study area) regarding extrapolation of type 1 (a) and type 2 (b). Extrapolation is expressed in percentage of the total area (log+1 transformed) for type 1 and type 2 novelty (Mesgaran et al., 2014). Type 1 novelty indicates areas in the projection space with at least one covariate being outside the univariate range of calibration data. Type 2 novelty indicates areas that are within the univariate range of calibration data but represent non-analogous covariate combinations. The spatial extrapolation was assessed across 500 species (average of replicates) under eight pseudo-absences sampling strategies (blue = generic; red = species-specific) when the calibration dataset included the original thinned set of occurrences (“Without adding spatial bias”) or a spatially biased subset of occurrences (“With adding spatial bias”), and when the set of pseudo-absences did not include (“Without background strata”) or included (“With background strata”) an environmentally stratified background of pseudo-absences. Grey numbers on the Y axis show percentage values without log+1 transformation.

Adding spatial bias to occurrences of the calibration data increased model extrapolation into novel covariate space for both NT1 and NT2 (Fig. 4, S6 and S7), but particularly for density specific (median value: ΔNT1 = +10.47%; ΔNT2 = +3.65%), target specific (ΔNT1 = +15.89%; ΔNT2 = +3.43%) and geographic specific (ΔNT1 = +1.30%; ΔNT2 = +0.03%) pseudo-absence strategies. Overall, adding a background sampling reduced the level of model extrapolation for all pseudo-absence strategies irrespective of adding spatial bias to occurrences (max values: NT1 < 0.25%; NT2 <0.01%; Fig. 4, S6 and S7).

## Discussion

We investigated how SDM predictions are influenced by pseudo-absence strategies in a study using high spatial resolution in observations and predictors. Our study moves beyond existing comparisons by investigating the performance of generic vs. species-specific pseudo-absence strategies in species-rich datasets that pose particular computational burdens. Using a comprehensive independent dataset of presence and absence records in the model evaluations, we found that both generic and species-specific strategies can yield high SDM performance. In particular, random, generic density, and geographic-specific pseudo-absence strategies provided best performance scores. Adding spatial bias to occurrence records decreased overall model accuracy, but did not challenge the general conclusions. Importantly, our study highlights potential extrapolation issues in pseudo-absence strategies, and we advocate to generally add a set of environmentally stratified pseudo-absences to cover the whole environmental space in model calibrations. This strategy may only be problematic in cases of strong and environmentally structured sampling biases. Given the overall marginal differences in performance observed between investigated pseudo-absence strategies the use of complex and computationally demanding pseudo-absence sampling approaches (e.g., target, density, geographic specific) may not always be justified compared to simpler approaches (e.g., random). Together, our study provides a better understanding of the potential effects of pseudo-absence sampling strategies on model performances in complex terrain.

### Selecting the best pseudo-absence strategy

By using a large and comprehensive set of generic and species-specific pseudo-absence strategies, we show that random, generic density, and geographic-specific provided overall best SDM performances in our topographically complex study area. Differences in the spatial patterns of pseudo-absence sets are obvious (e.g., *Androsace helvetica* in Fig. 1) and support why previous studies found different model outcomes between various pseudo-absence strategies (Chefaoui & Lobo, 2008; Hanberry et al., 2012; Wisz & Guisan, 2009). Our results are overall in line with assessments reporting higher performance of SDMs when pseudo-absences are selected randomly or around occurrences of the species (Barbet-Massin et al., 2012; F. Graf et al., 2006; Kramer-Schadt et al., 2013; Wisz & Guisan, 2009), yielding more constrained spatial projections of suitable habitat (Chefaoui & Lobo, 2008; Hanberry et al., 2012). While generic approaches (random, environmentally proportional- or equal-stratified, target group or density dependent) are frequently documented, species-specific approaches have received little attention so far, especially when working with species-rich datasets.

Scientists attempting to model the distribution of many species at the same time may favour one unique generic pseudo-absence approach over species-specific ones. This will reduce computational burdens and complexity of fine-tuning the pseudo-absence selection for each species individually. The differences in SDM performances observed between the best approaches also supports that a generic random sampling approach might be more appropriate to model large species sets in the framework of our study. Hence, the random sampling approach has the benefit of being set up easily, rapidly and applied to each species in the same way, and to cover a sufficient portion of the environmental space in model calibrations to avoid potential spatial extrapolation issues. Therefore, the use of complex and computationally demanding pseudo-absence approaches (e.g., target, density, geographic specific) may not always be justified compared to the simpler random approach. Our study provides guidelines and the first overview of the performance of species-specific and generic pseudo-absence strategies in SDMs and in species-rich datasets that pose particular computational challenges.

### Effect of increased sampling bias

Increasing spatial bias in the occurrence records decreased the overall model accuracy, but did not result in different conclusions concerning the choice of the best approaches. The target specific and density specific approaches were particularly affected by the added sampling bias as they showed the highest drop in model performances (Fig. 3c-d). Overall, we found that random, generic density, and geographic specific strategies performed best, and that the geographic specific strategy remained the best approach when a background of stratified pseudo-absences was added. Both generic density and geographic specific approaches have higher densities of pseudo-absences collected in regions more frequently sampled, and can therefore compensate for some level of sampling bias (Hertzog et al., 2014; Kramer-Schadt et al., 2013). Surprisingly, and in contrast to recommendation of previous studies (Elith & Leathwick, 2007; Mateo et al., 2010; Phillips et al., 2009), both target group sampling strategies (generic and species-specific) showed lower performance. This might be due to the opportunistic nature of the presence data in our data where observers did not necessarily search for specific target groups or checklists. Also, the lower performance may be related to the lower environmental coverage of the calibration data of the target group strategies which only sample cells that have at least one record. Hence, high levels of spatial extrapolation were found within environmental domains not covered by the calibration data, especially when no background of stratified pseudo-absences were included, as was also found by Iturbide et al. (2015). Our results are also in line with studies recommending the random approach, if no suitable set of species may be chosen as a representative target-group (Cerasoli et al., 2017). Hence, the selection of one specific target group (e.g., a clade or family, or a functional group) may considerably affect model predictions (Righetti et al., 2019). The use of the target specific strategy will especially be challenging for studies trying to model a very large set of species, since this strategy needs to be adapted to each single species. As suggested by Stokland et al. (2011), a random sampling strategy is more transparent and rests upon fewer assumptions. Our study stresses the importance of well-distributed occurrence and absence data across environmental and geographic space as a prerequisite for building sound SDMs (Lobo & Tognelli, 2011). Overall, the geographic-specific pseudo-absence strategy is a valuable species-specific approach for compensating spatial bias in species records, but its complexity and high computational cost make it a challenging strategy to implement for species-rich datasets.

Spatial thinning of occurrences might be an adequate alternative for bias compensation when occurrence datasets are spatially biased and when enough data are available (Kramer-Schadt et al., 2013). In our study, occurrences were thinned by 300 m because it significantly reduced spatial autocorrelation in model residuals, which is one of the prerequisites of applying sound statistical models. The spatial thinning is particularly relevant at high spatial resolution because it allows to reduce pseudo-replication of the same plant population, and therefore reduces risks of overfitting the model to specific environmental conditions. However, the spatial thinning does not solve the problem of unsampled environments. By adding pseudo-absences into unsampled environments, we assume the species to be absent at this location. On the contrary, when pseudo-absences are not added to unsampled environments, we might allow the model to predict the presence of the species at this location even if it is in fact absent at this location. Therefore, the selection of a pseudo-absence strategy, especially when high spatial bias in occurrences is expected, should be performed with caution and also consider the goals of the project and for which purpose the modelled outputs will be used. Should we risk predicting presence of a species in unsuitable areas (risk of potential overprediction due to erroneous extrapolation in case of truncated niches) or should we assume that the distribution of a species is limited by the conditions under which it has been observed (risk of potential underprediction due to avoiding extrapolation)? This might be particularly problematic if SDM-based predictions are used to define suitable areas for conservation purposes or species re-introductions, because model overpredictions might then introduce incorrect guidance to field biologists, environmental practitioners and stakeholders. Models from biased samples should therefore always be checked against expert knowledge on the species’ ecology.

Finally, the spatial bias that we introduced in our datasets overemphasized the most easily accessible and famous botanical areas of Switzerland. The environmental coverage of the calibration data, in turn, decreased for all species-specific strategies, resulting in higher levels of spatial extrapolation when spatial bias was added (Fig. 4a-b). In contrast, generic strategies were not much affected by the inclusion of spatial bias and the ranking of the best strategies remained almost unchanged. Compared to other studies which often do not use spatial thinning, the spatial thinning of occurrences that we applied already strongly reduced the spatial bias in our dataset by reducing the number of occurrences from heavily sampled regions. In addition, the bias that we introduced might be too low or the results we obtained are specific to the high topographic complexity of our study area. This potentially explains why commonly reported sampling strategies, such as target-group (e.g., Phillips et al., 2009), often applied in studies performed at coarser resolutions, do not perform well in our case and why the random strategy seems to be more performant in our study. Overall, further research is needed to determine how sampling bias can be compensated by means of different pseudo-absence strategies, at different spatial resolution (>5 km to <100 m) and ecological context (complex terrain versus smooth ecological gradients).

### Including an environmentally stratified background of pseudo-absences

Incomplete environmental niche representation in calibration datasets of niche-based models can lead to highly uncertain extrapolation into new environments not covered by the calibration dataset (Mesgaran et al., 2014; Thuiller et al., 2004, Zurell et al. 2012). In our study, we found considerable risks of extrapolations into novel covariate space for target, density and all species-specific pseudo-absence strategies. This mostly concerned alpine regions or lakes due to the low representation of high elevation sites and lake borders in the calibration data. Adding an environmentally stratified background of pseudo-absences enabled us to strongly decrease spatial extrapolation (Type 1 and 2) into new environmental space, and clearly improved model performance of species-specific strategies. Furthermore, such background remains essential if the calibration dataset covers only portions of the environmental space of regions the models are projected to, which often happens if models are projected into current or future environmental conditions (Thuiller et al., 2004). However, while the stratified background primarily adds constraints to environments in which no species should occur (lakes, high mountains), it may also add absences to unsampled environments in which species actually occur. In the latter case, we enforce truncated niches which can turn out to be problematic for projections under climate change, as we then potentially underpredict the species’ potential distribution. Therefore, studies aiming at predicting the potential distribution of species under climate change, should also, whenever possible, consider future potential environmental conditions by means of larger calibration areas by adding historical or fossil records (Descombes et al., 2015; Maiorano et al., 2013; Thuiller et al., 2004) or by predicting species only into areas where similar environmental conditions are observed in the calibration dataset (safe projections). While SDMs calibrated on only a fraction of the environmental space might lead to unrealistic overpredictions of species distribution towards environmental extremes (Thuiller et al., 2004), including a set of environmentally stratified pseudo-absences enables to avoid unpredictable effects on the tails of the species response curves.

### Recommendations and conclusions

Based on our results and the considerations discussed above, we formulate four recommendations for selecting pseudo-absences for building SDMs (see Table 1):

**Table 1.**
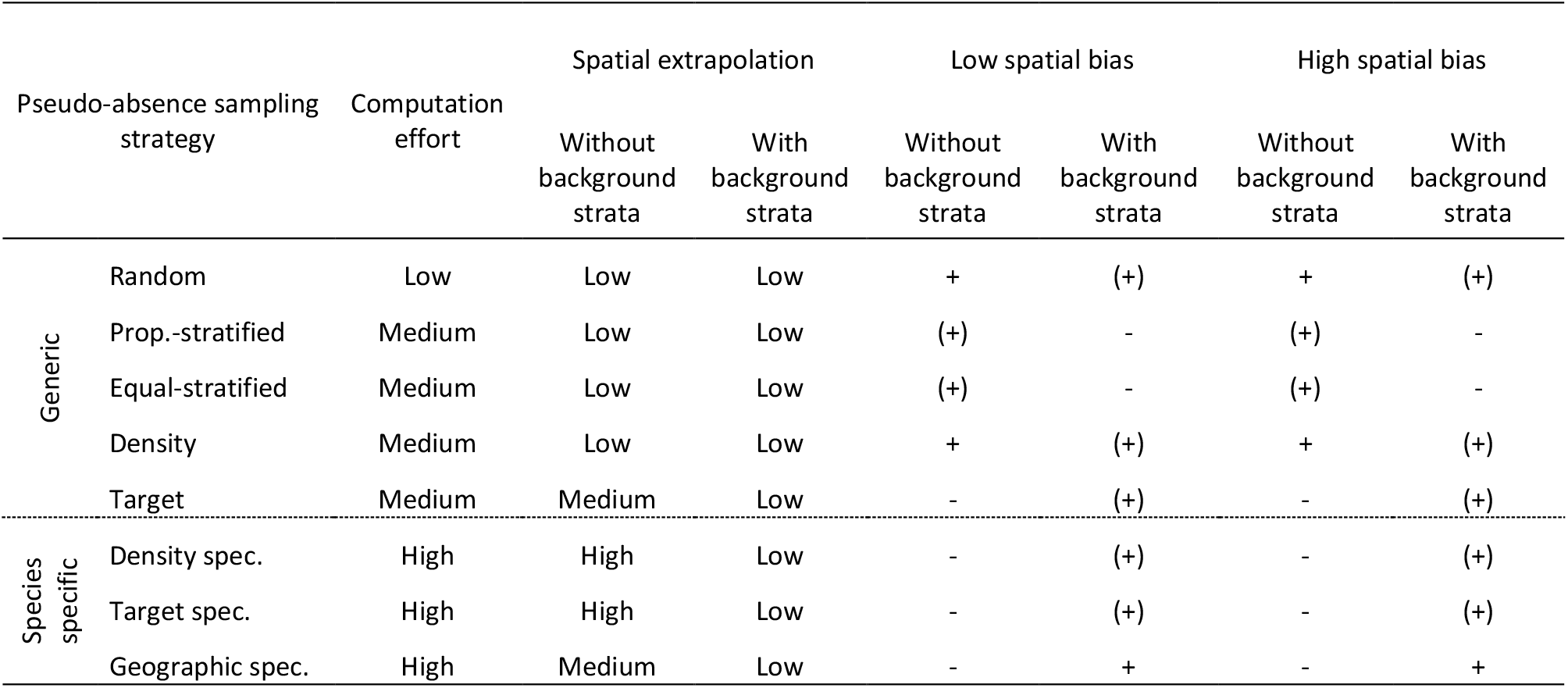
Recommendation of pseudo-absence sampling strategies (generic and species-specific) to be used if low or high spatial bias in datasets is detected or assumed, and if a set of environmentally stratified background of pseudo-absences is made available to be included or not. Computational effort represents a relative measure of the time necessary to prepare and compute the pseudo-absence set on a “per species” basis (Low = up to few seconds, Medium = few minutes, High = >5 minutes). The risk of spatial extrapolation from the calibration data in the projection space regarding spatial extrapolation of type 1 and type 2 is summarized based on our results. + = recommended; (+) = recommended with caution, - = not recommended

#### 1. Spatial thinning of occurrences

If sufficient observations are available, occurrences should be spatially thinned with a minimal distance of the grid resolution, and ideally to a horizontal grid resolution that is ecologically informed for the taxonomic group of interest (e.g., by the average spatial extent of populations). Such spatial thinning, particularly at high spatial resolution, reduces spatial auto-correlation and pseudo-replication of observations belonging to the same population. As a result, it reduces the risk of overfitting SDMs to specific environmental conditions.

#### 2. Extrapolation into novel environments

In datasets with strong environmental gradients, many pseudo-absence strategies may lead to unconstrained response curves that create extrapolation problems. We thus advocate that pseudo-absence strategies should include a minimal set of environmentally stratified pseudo-absences covering the complete range of observed environmental conditions in the study area. Our results suggest that including such a set helps to avoid unrealistic overpredictions of species towards environmental gradient extremes, especially for target, density and all species-specific pseudo-absence sampling strategies. If SDMs are transferred to different study areas or time periods (future/past environmental conditions), calibration data should include the same environmental space as the projection region (adding historical or fossil records), or projections should only be performed into areas where similar environmental conditions are observed as in the calibration dataset (safe projections).

#### 3. Choosing the best pseudo-absence strategy

We recommend the use of the random, generic density, or geographic specific pseudo-absence strategy. Given the high computational demand of species-specific approaches, their use may not always pay off in species rich datasets.

#### 4. Spatially biased occurrence datasets

For spatially biased occurrence datasets, we recommend to use a generic density approach or a geographic specific approach, ideally with additionally including a minimum set of environmentally stratified pseudo-absences outside the observed environment as defined by the environmental predictors. This helps to avoid spatial extrapolation into unsampled environmental space due to unconstrained niches towards the edges of the observed environment. Note that the spatial bias is often removed or severely reduced by a spatial thinning of the occurrences.

In summary, our findings pave the way towards a better and more comprehensive assessment of the effects of pseudo-absence strategies on SDM performance. We illustrate the effects of diverse strategies and we conclude with recommendations. Although the choice of the best strategy depends on several criteria (spatial bias, spatial extent and resolution, environmental complexity, number of species to model), studies should carefully weigh potential SDM performance increase against computational demand when using complex, species-specific pseudo-absence strategies (e.g., target, density, geographic specific). Our results provide valuable additions to the debate on the choice of pseudo-absence strategies in SDMs. Further research might investigate the performance of an ensemble approach by averaging results of different pseudo-absence strategies. Finally, while many studies rely on opportunistically sampled data from various sources (e.g., museum datasets, citizen science programs) often gathered without adequate sampling designs, we recommend citizen science databases (e.g., Gbif, iNaturalist, InfoFlora) to move towards incorporating stratified sampling designs allowing the collection of true absences by citizen scientists.

## Acknowledgements

We thank InfoFlora and the Swiss Biodiversity Monitoring programme who kindly provided the plant occurrence data, and Tobias Roth and Michael Jutzi for help with data extraction. We also thank Christian Ginzler for the help with generating the forest environmental predictor layers. This study was supported by the Swiss Data Science Center (SDSC) grant no. c17-07 (SPEEDMIND) project to NEZ, DZ, PD and DNK. DNK. & NEZ acknowledge funding from: The WSL internal grant exCHELSA, and ClimEx, the 2019-2020 BiodivERsA joint call for research proposal under the BiodivClim ERA-Net COFUND programme (project ‘FeedBaCks’) with the national funder Swiss National Science Foundation (20BD21_193907). DNK acknowledges funding to the ERA-Net BiodivERsA - Belmont Forum, with the national funder Swiss National Foundation (20BD21_184131), part of the 2018 Joint call BiodivERsA-Belmont Forum call (project ‘FutureWeb’). NEZ, PB and YC acknowledge additional funding to Origin.Alps from the Swiss National Science Foundation (310030L_170059). DZ acknowledges funding from the German Science Foundation (ZU 361/1-1).

## Data availability

We provide an R package containing functions to generate pseudo-absence points based on the approaches presented in this study on Githubb (https://github.com/filBe87/PseuAbs).

## Authors’ contributions

PD conceived the ideas with the help of YC, NEZ, DZ, DNK, DR and ROW. PD and ROW collected and prepared the data. PD performed the analyses. PB cleaned and prepared the pseudo-absences functions made available through this manuscript. PD led the writing of the manuscript. All authors contributed critically to the drafts and gave final approval for publication.

