## Supplementary materials for "Strategies for sampling pseudo-absences for species distribution models in complex mountainous terrain"

**Note S1**

*Spatial thinning of occurrences*

We investigated to what degree spatial thinning of species occurrences reduces spatial autocorrelation in model residuals, by removing occurrences of the 500 selected plant species at increasing distances from 0 m to 5000 m (steps of 25 m from 0 to 600 m, of 50 m from 600 to 800 m, of 100 m from 800 to 1000 m, and of 500 m from 1000 to 5000 m). Our investigations were based on both generalized linear models (glm; McCullagh, 1983) and generalized additive models (gam; Hastie & Tibshirani, 1987; Guisan, Edwards, & Hastie, 2002). To this end, we selected a set of 10'000 pseudo-absences randomly across Switzerland, and we repeated this selection three times. For each species, we then related occurrences and pseudo-absences to 10 environmental predictors (see the environmental predictors section in the main text). All selected environmental predictors showed pairwise Pearson correlations $|r| < 0.6$ (Dormann et al., 2013). Models were built with each of the three replicates of pseudo-absences and equal weights were given for presences and pseudo-absences in the modelling process (i.e. the weighted sum of presence equals the weighted sum of pseudo-absence; Barbet-Massin, Jiguet, Albert, & Thuiller, 2012). We used the Moran's I test from the *ape* R package (Paradis & Schliep, 2019) to assess spatial autocorrelation in model residuals and averaged values across replicates. The optimal thinning distance was identified such that 95% of the highest Moran's I values for the corresponding thinning distance were below the 5% of the lowest Moran's I when no thinning distance was applied (i.e. thinning distance = 0). To assess the effect of the spatial thinning of occurrences on model accuracies (Fig. S2), we evaluated the models based on a five-fold repeated split-sample test (training set = 80%, evaluation set = 20%) by using the True Skill Statistic (Allouche, Tsoar, & Kadmon, 2006). TSS evaluates the ability of the model to discriminate presences from absences, and scales between -1 and 1 (0 indicating random predictions). Predictive performances of models with TSS > 0.4 are considered as reliable (i.e. excellent TSS > 0.75; good 0.40 < TSS < 0.75; poor TSS < 0.40) (Landis & Koch, 1977).

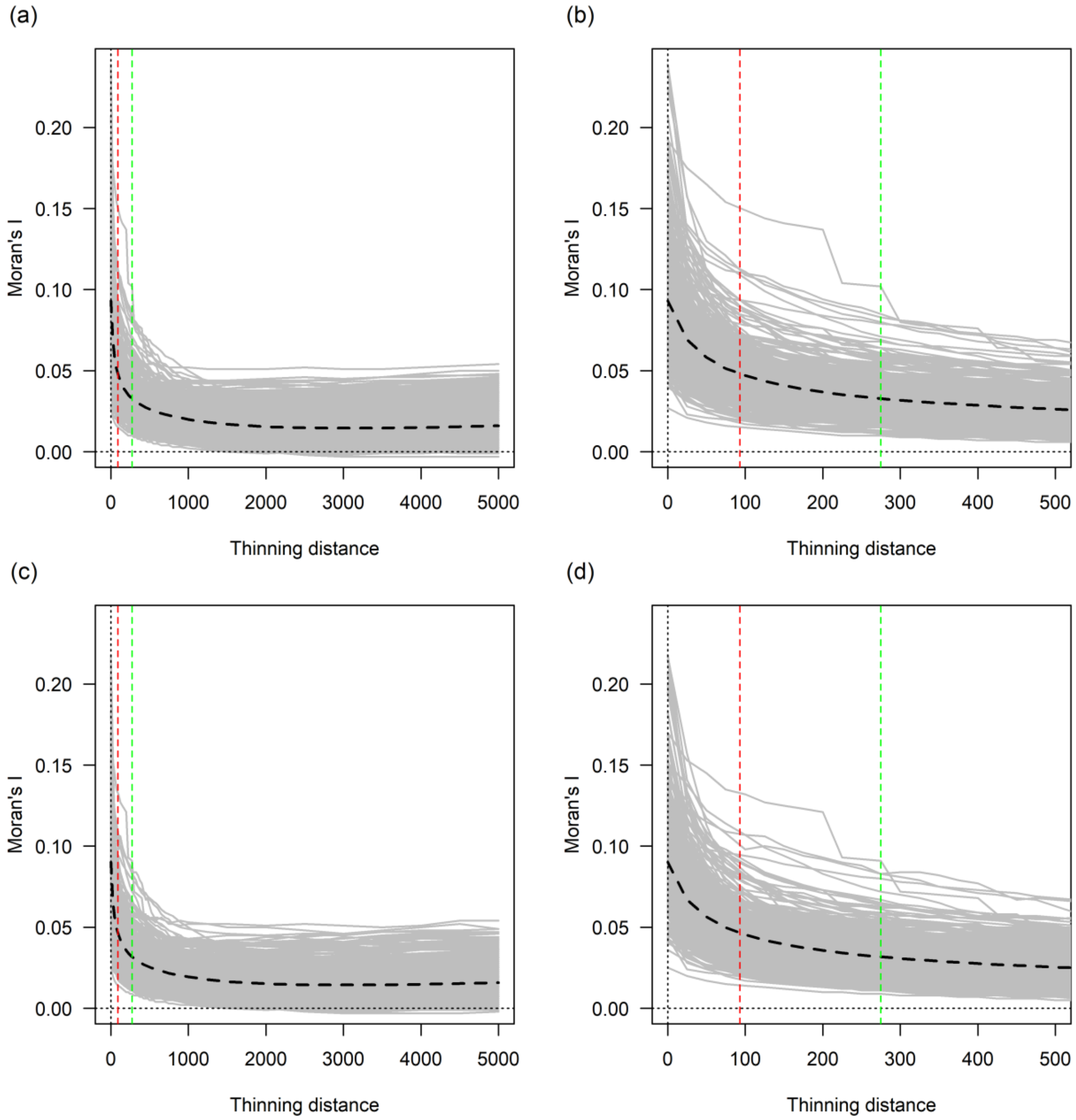


**Figure S1.** Spatial autocorrelation in model residuals (Moran's I) from glm (a, b) and gam (c, d) models in relation to the distance used to thin the original occurrence data. Left (a, c) and right (b, d) panels differ in the represented range of thinning distances. Grey lines represent Moran's I values of the 500 species studied (averaged across the 3 replicates) and dotted black lines the average across all species. Dotted red lines show the distance corresponding to the predictor resolution (93 m). Dotted green lines show the optimal thinning distance (275 m here).

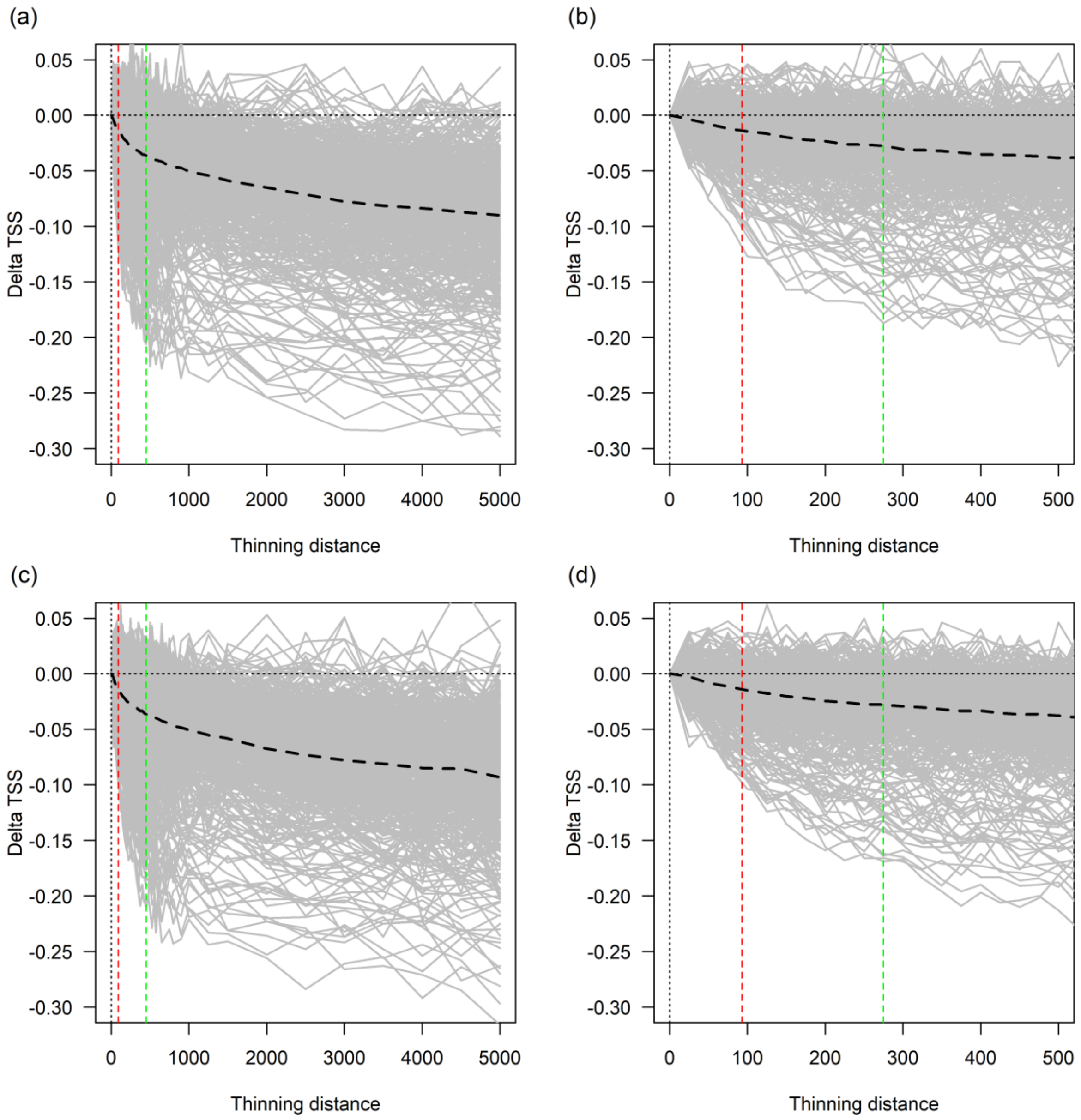


**Figure S2.** Effect of the spatial thinning of species occurrences on TSS-based (split sampling derived) model prediction accuracy in glm (a, b) and gam (c, d) models, expressed as the TSS difference between models calibrated with and without spatial thinning. Left (a, c) and right (b, d) panels differ in the represented range of thinning distances. Grey lines represent TSS values of the 500 selected species (averaged across the 3 replicates) and dotted black lines the average across all species. Dotted red lines show the distance corresponding to the predictor resolution (93 m). Dotted green lines show the optimal thinning distance (275 m here).

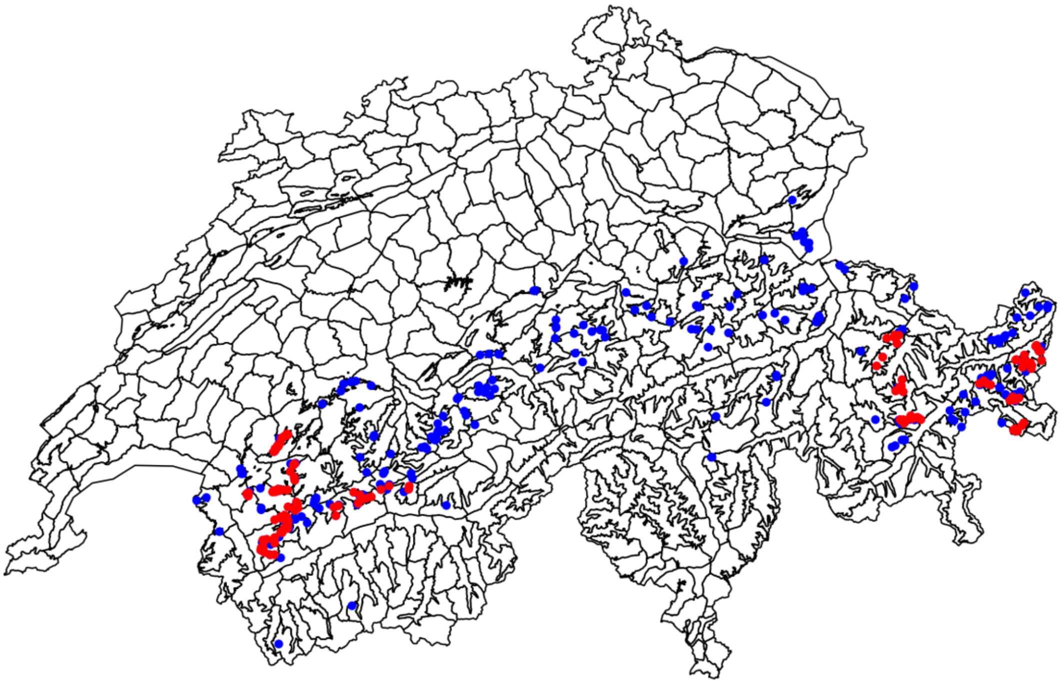

**Figure S3.** Map representing the polygons used in the Welten & Sutter (1982) plant distribution atlas of Switzerland, defined by experts and based on political and watershed boundaries, and the treeline. This polygon dataset was originally used to map the Swiss Flora, and was used here to add sampling bias to the occurrence data. For every species, spatial bias was introduced by selecting 150 occurrences from the 10% most sampled Welten & Sutter polygons. The map shows the example of *Androsace helvetica* (L.) All., where all coloured points represent the original occurrence data of the species (n = 388), and red points (n = 150) the occurrences of the spatially biased subset.

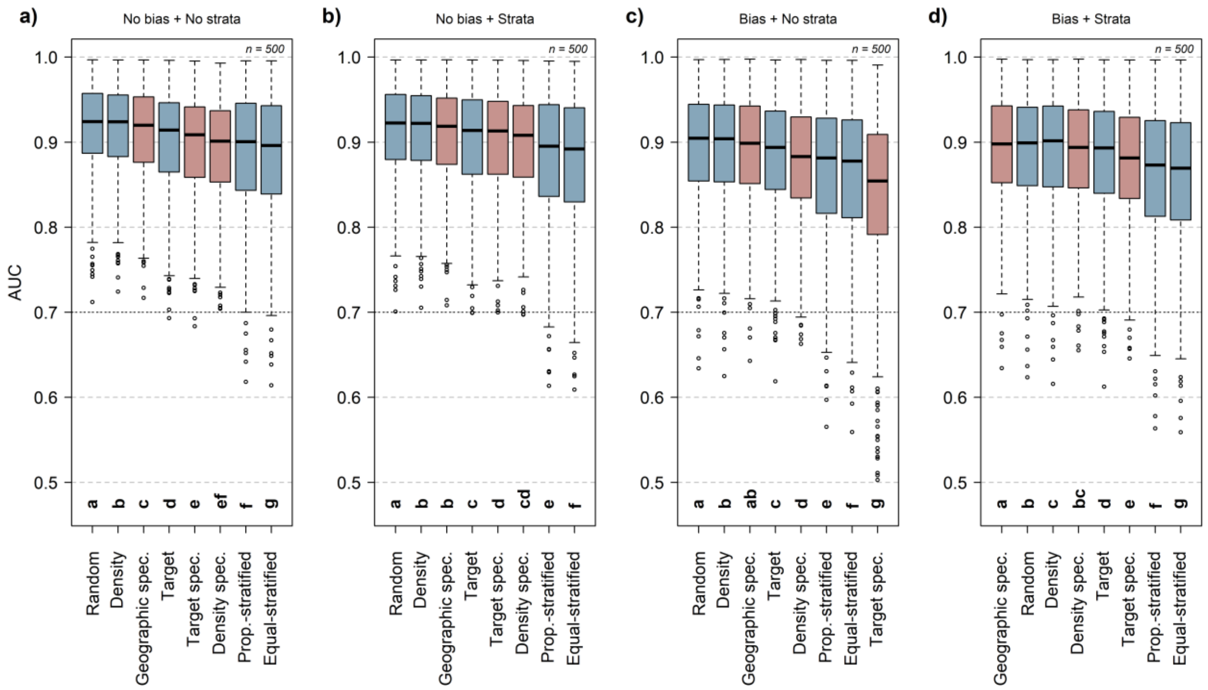


**Figure S4.** SDM performance (AUC metric) of ensemble models assessed for 500 plant species (average of replicates) under eight pseudo-absences sampling strategies (blue = generic; red = species-specific). Models were run with the original thinned set of occurrences (a, b; "No bias") or with a spatially biased subset of occurrences (c, d; "Bias"). Pseudo-absences were sampled following the eight pseudo-absences sampling strategies and included (b, d; "Strata") or did not include (a, c; "No strata") an environmentally stratified background of pseudo-absences in the calibration dataset. The pseudo-absence sampling strategies are ranked according to their average performances (high to low from left to right). The dotted black line represents the thresholds above which models are considered to have reliable performances (AUC > 0.7). All pairwise comparisons were run with paired Wilcox tests with Bonferonni correction for multiple tests and displayed following a letter-based representation significant difference (p-value > .05).

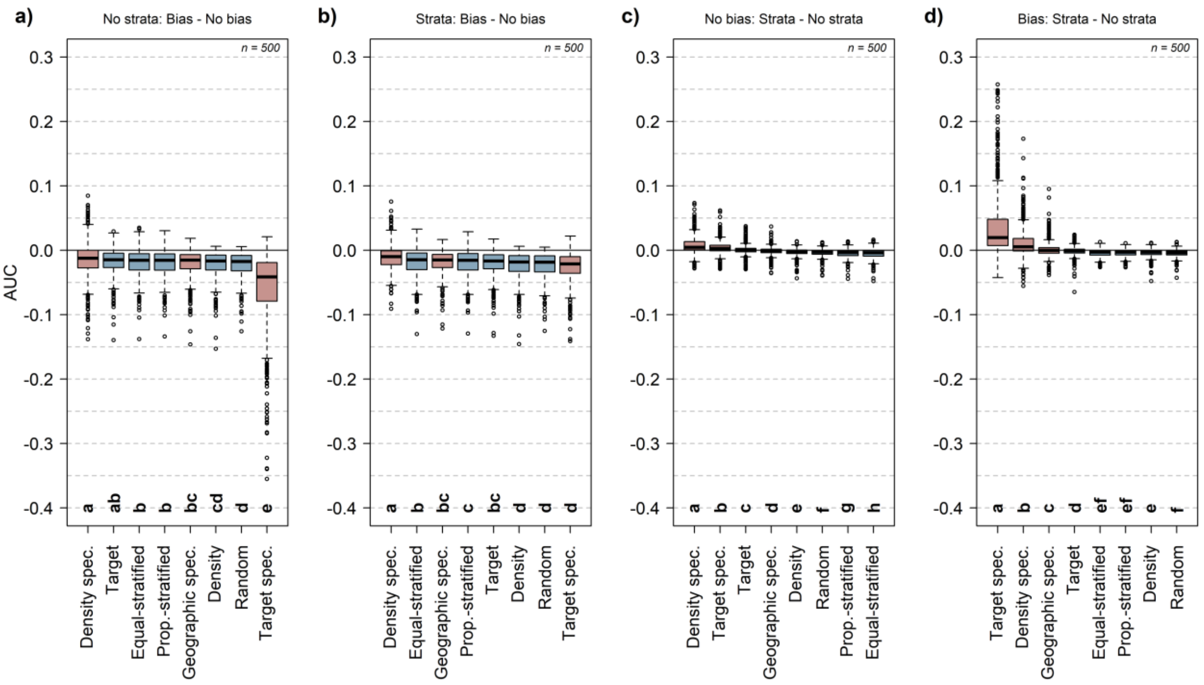


**Figure S5.** Differences in SDM performance (AUC metric) of ensemble models assessed for 500 plant species (average of replicates) under eight pseudo-absences sampling strategies (blue = generic; red = species-specific). *Panel a* and *b* represent the effect of spatial bias added in the set of occurrences on the model performance ("Bias – No bias") when the set of pseudo-absences did not include (a; "No strata") or included (b; "Strata") an environmentally stratified background of pseudo-absences in the calibration dataset. *Panel c* and *d* represent the effect of including an environmentally stratified background of pseudo-absences in the pseudo-absences sets ("Strata – No strata") when the models were calibrated with the original thinned set of occurrences (c; "No bias") or with a spatially biased subset of occurrences (d; "Bias"). The pseudo-absence sampling strategies are ranked according to their average difference in performances (high to low from left to right). All pairwise comparisons were run with paired Wilcox tests with Bonferonni correction for multiple tests and displayed following a letter-based representation of significant difference (p-value > .05).

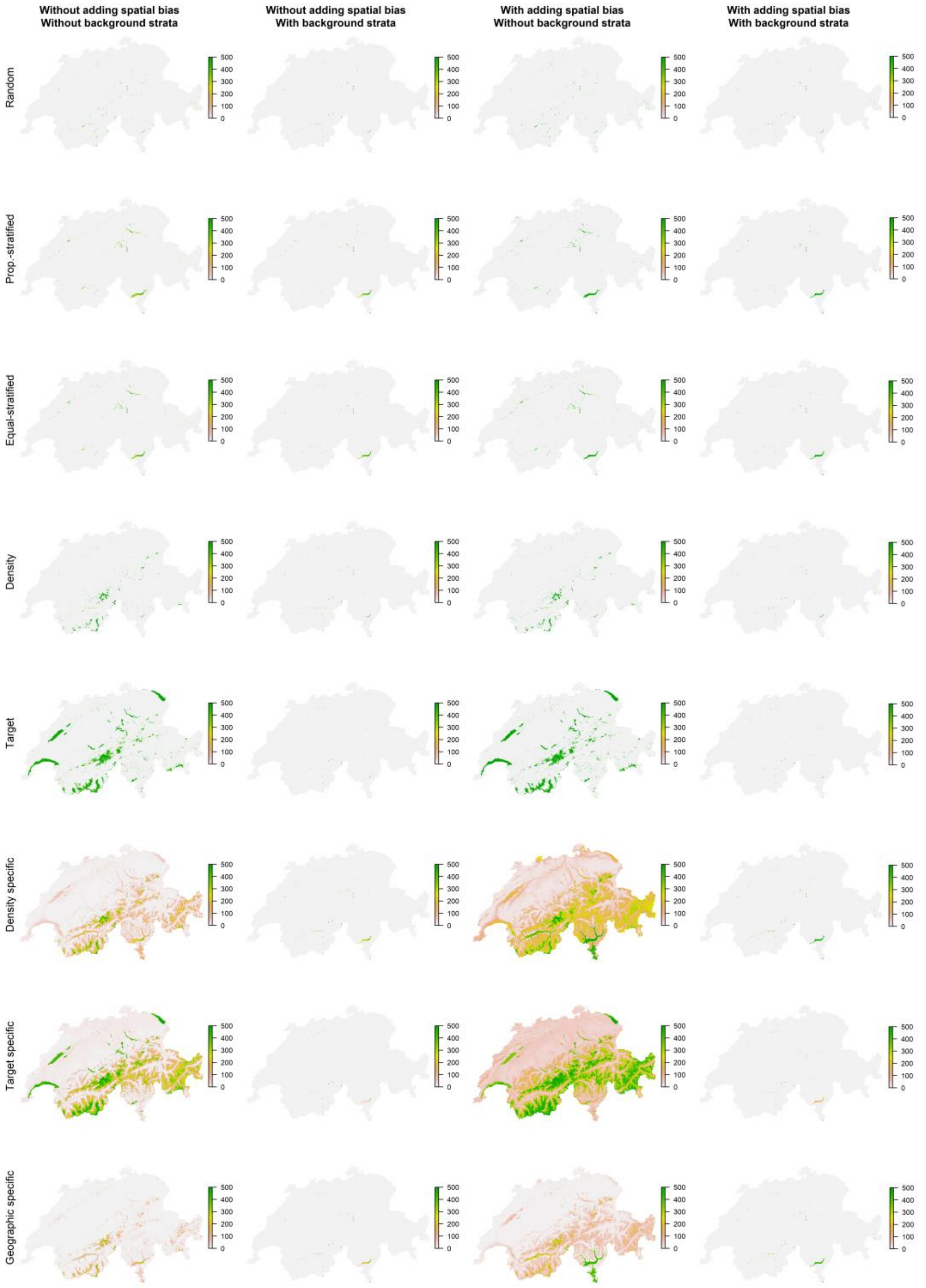


**Figure S6.** Quantification of the level of extrapolated projection area of type 1 novelty across the 500 plant species for eight pseudo-absence sampling strategies, with or without adding a background of environmentally stratified pseudo-absences, and with or without adding spatial bias in occurrences. For each species, we retained cells presenting type 1 novelty values (NT1; values < 0) and summed this information across all species. Type 1 novelty values corresponds to extrapolation outside the univariate range of at least one of the covariates compared to the reference data (Mesgaran, Cousens, & Webber, 2014). The coloured scale represents the number of species presenting type 1 novelty at a given location.

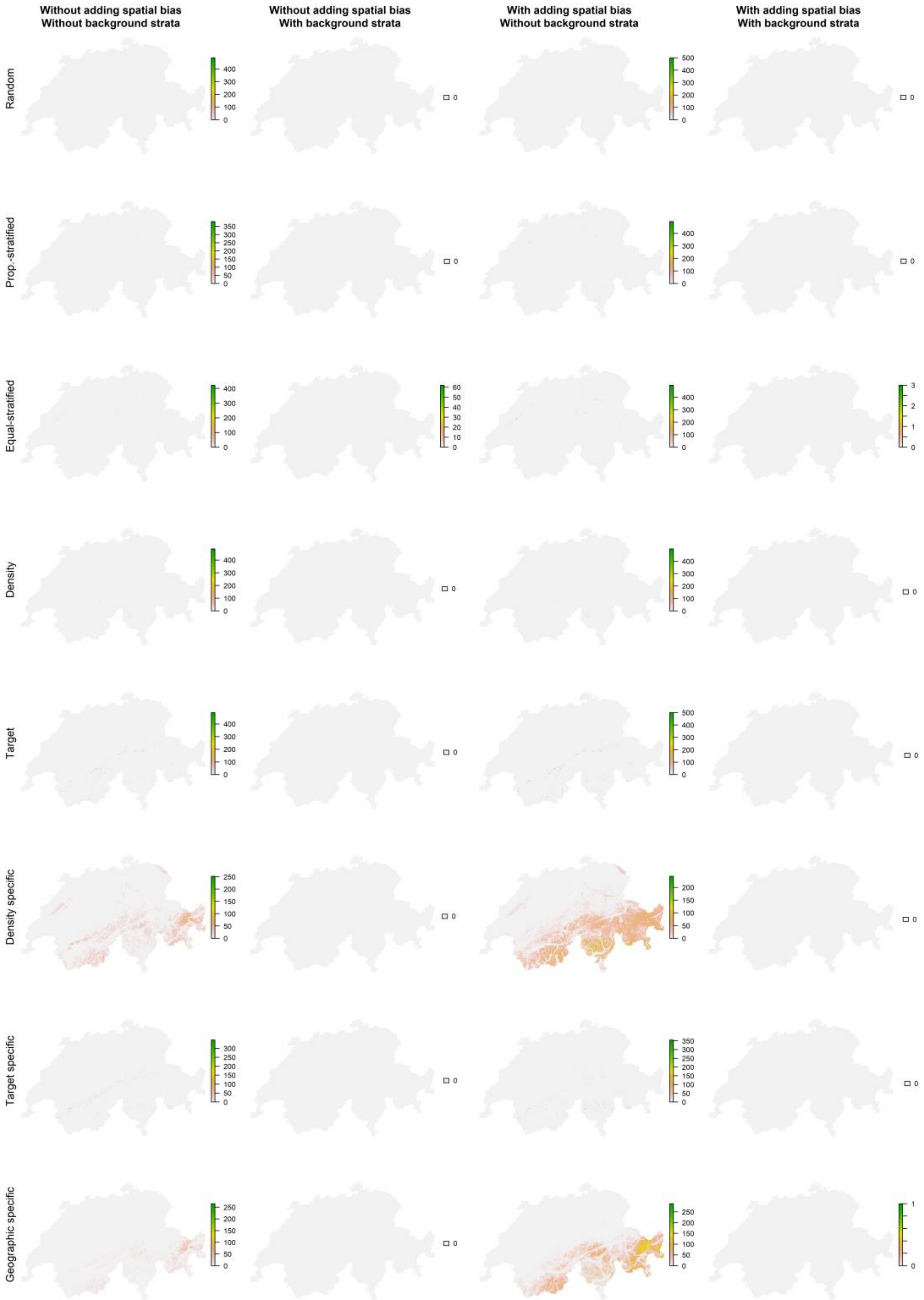


**Figure S7.** Quantification of the level of extrapolated projection area of type 2 novelty across the 500 plant species for eight pseudo-absence sampling strategies, with or without adding a background of environmentally stratified pseudo-absences, and with or without adding spatial bias in occurrences. For each species, we retained cells presenting type 2 novelty values (NT2; values > 1) and summed this information across all species. Type 2 novelty corresponds to novel covariate combinations which are within the univariate range of covariates compared to the reference data (Mesgaran et al., 2014). The coloured scale represents the number of species presenting type 2 novelty at a given location.

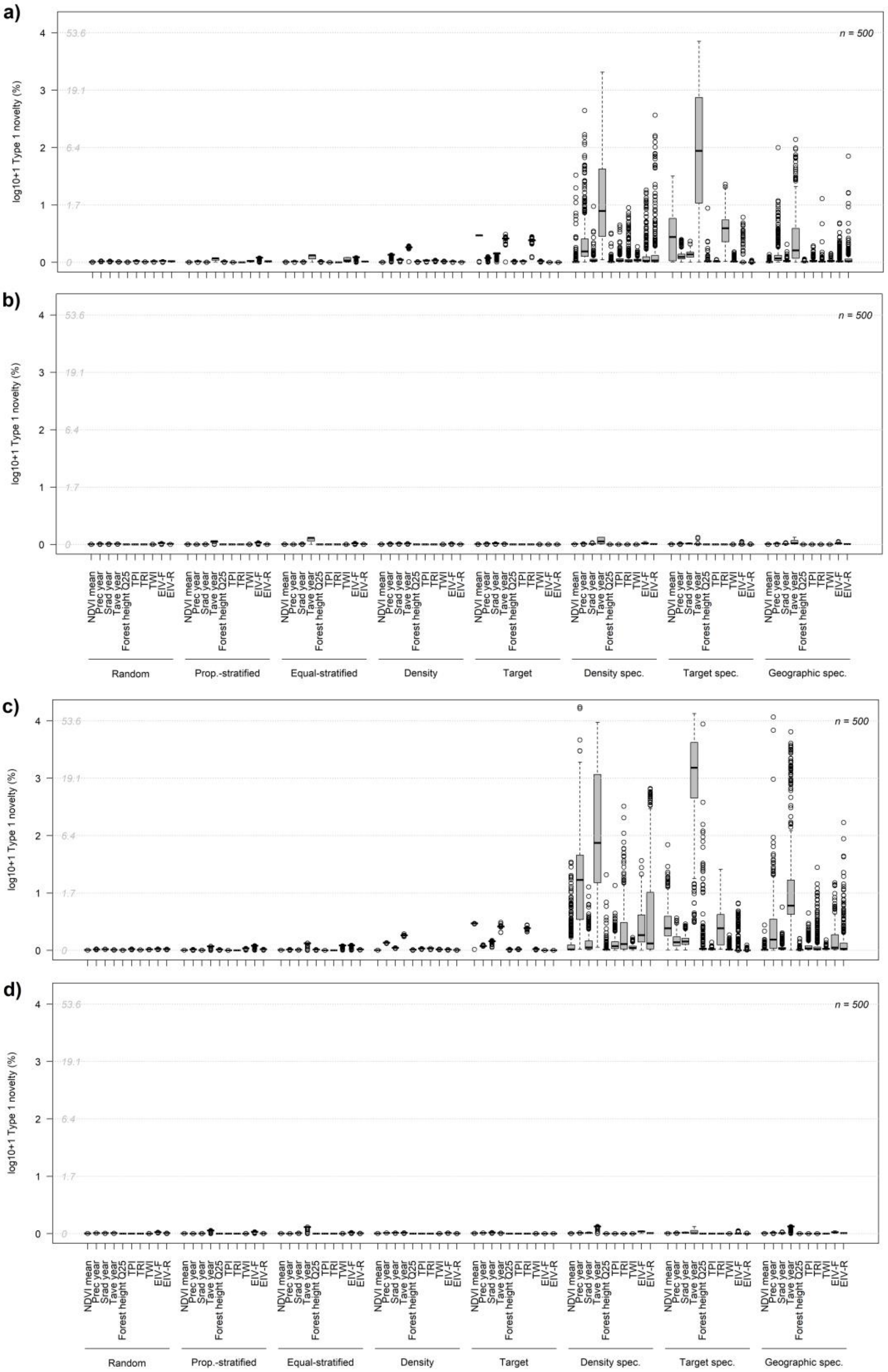


**Figure S8.** Contribution of each predictor to type 1 novelty across the 500 plant species expressed as percentage of the total area (log+1 transformed) for eight pseudo-absence sampling strategies without (a, c) or with (b, d) adding a background of environmentally stratified pseudo-absences and without (a, b) or with (c, d) adding spatial bias to occurrences. Grey numbers on the Y axis show percentage values without log+1. transformation.

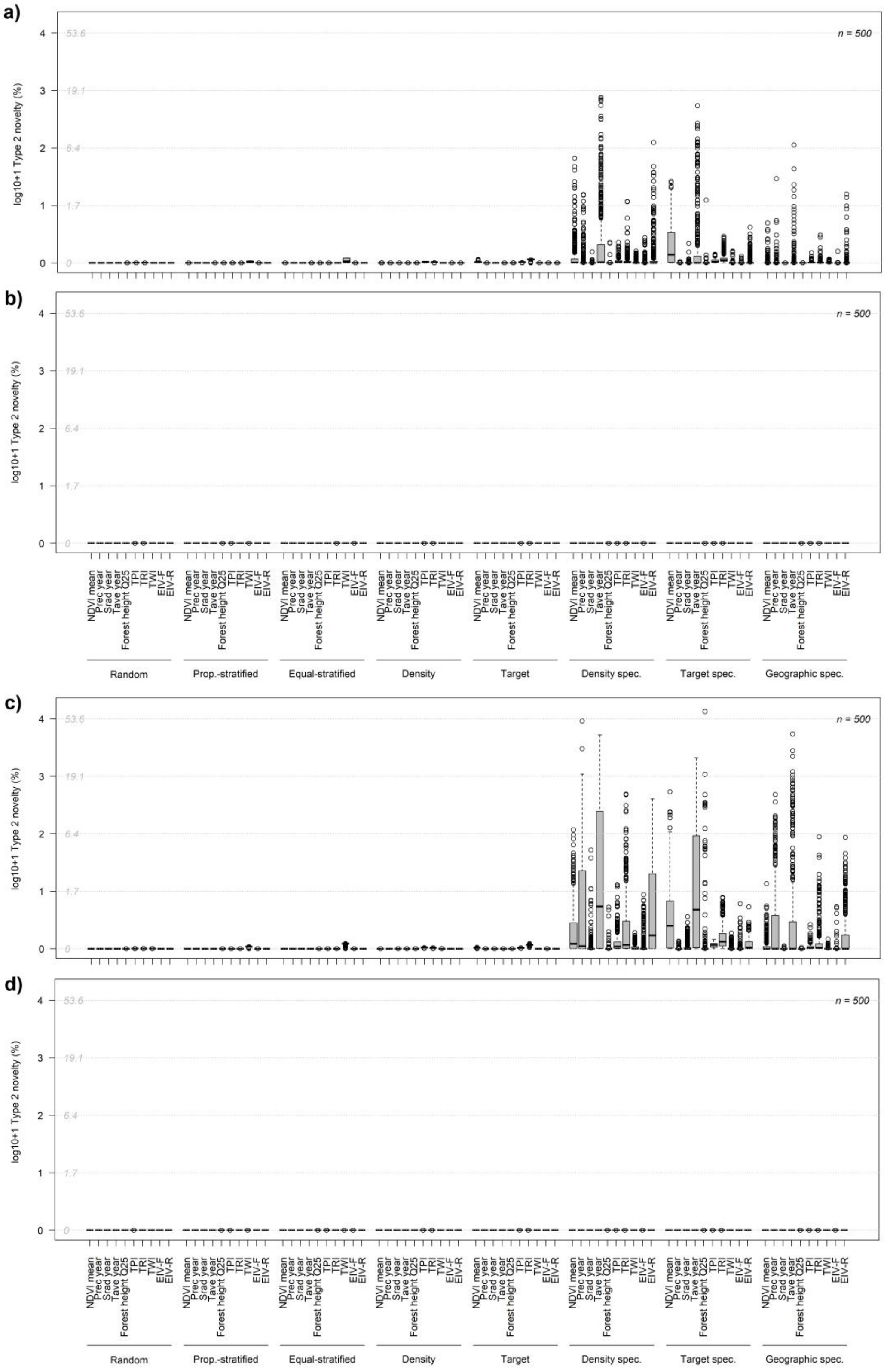


**Figure S9.** Contribution of each predictor to type 2 novelty across the 500 plant species expressed as percentage of the total area (log+1 transformed) for eight pseudo-absence sampling strategies without (a, c) or with (b, d) adding a background of environmentally stratified pseudo-absences and without (a, b) or with (c, d) adding spatial bias to occurrences. Grey numbers on the Y axis show percentage values without log+1. transformation.

**Table S1.** SDM predictive performance (TSS metric, evaluated on the independent presence/absence dataset) of ensemble models assessed across 500 plant species under eight pseudo-absences sampling strategies (average among pseudo-absence replicates) for the ensemble and single models of five algorithms (glm, gam, gbm, rdf and max). The three best evaluations are highlighted in bold and the best pseudo-absences sampling strategies are ranked (Rank). Glm = generalized linear models, gam = generalized additive model, gbm = gradient boosting machines, rdf = Random Forest, max = maximum entropy.

| Pseudo-absence selection strategy | | | Ensemble evaluation (TSS) | Single model evaluations (TSS) | | | | | | Rank |
|---|---|---|---|---|---|---|---|---|---|---|
| | | | | mean | glm | gam | gbm | rdf | max | |
| Without adding spatial bias | Without background strata | Random | **0.710** | **0.698** | **0.687** | **0.695** | **0.687** | **0.722** | **0.697** | 1 |
| | | Prop.-stratified | 0.650 | 0.639 | 0.633 | 0.644 | 0.620 | 0.659 | 0.640 | 7 |
| | | Equal-stratified | 0.643 | 0.632 | 0.626 | 0.638 | 0.611 | 0.650 | 0.634 | 8 |
| | | Density | **0.706** | **0.693** | **0.682** | **0.690** | **0.685** | **0.717** | **0.693** | 2 |
| | | Target | 0.681 | 0.670 | 0.658 | 0.664 | 0.662 | 0.697 | 0.671 | 4 |
| | | Density spec. | 0.654 | 0.633 | 0.621 | 0.631 | 0.629 | 0.644 | 0.640 | 6 |
| | | Target spec. | 0.671 | 0.658 | 0.646 | 0.650 | 0.647 | 0.683 | 0.663 | 5 |
| | | Geographic spec. | **0.698** | **0.682** | **0.670** | **0.680** | **0.676** | **0.702** | **0.684** | 3 |
| | With background strata | Random | **0.701** | **0.689** | **0.676** | **0.686** | **0.679** | **0.716** | **0.686** | 1 |
| | | Prop.-stratified | 0.640 | 0.628 | 0.615 | 0.626 | 0.615 | 0.657 | 0.630 | 7 |
| | | Equal-stratified | 0.632 | 0.620 | 0.606 | 0.617 | 0.607 | 0.649 | 0.622 | 8 |
| | | Density | **0.700** | **0.687** | **0.674** | **0.684** | **0.678** | **0.713** | **0.683** | 2 |
| | | Target | 0.681 | 0.670 | 0.657 | 0.662 | 0.666 | 0.697 | 0.667 | 4 |
| | | Density spec. | 0.670 | 0.653 | 0.642 | 0.649 | 0.648 | 0.675 | 0.649 | 6 |
| | | Target spec. | 0.678 | 0.667 | 0.654 | 0.659 | 0.662 | 0.695 | 0.664 | 5 |
| | | Geographic spec. | **0.696** | **0.681** | **0.668** | **0.678** | **0.674** | **0.706** | **0.678** | 3 |
| With adding spatial bias | Without background strata | Random | **0.661** | **0.649** | **0.641** | **0.652** | **0.649** | **0.659** | **0.643** | 1 |
| | | Prop.-stratified | 0.612 | 0.600 | 0.603 | 0.608 | 0.592 | 0.596 | 0.603 | 6 |
| | | Equal-stratified | 0.605 | 0.594 | 0.598 | 0.603 | 0.584 | 0.588 | 0.598 | 7 |
| | | Density | **0.659** | **0.646** | **0.640** | **0.649** | **0.646** | **0.657** | **0.639** | 2 |
| | | Target | 0.643 | 0.630 | 0.625 | 0.632 | 0.631 | 0.639 | 0.622 | 4 |
| | | Density spec. | 0.628 | 0.603 | 0.587 | 0.619 | 0.606 | 0.596 | 0.607 | 5 |
| | | Target spec. | 0.568 | 0.551 | 0.539 | 0.534 | 0.560 | 0.559 | 0.563 | 8 |
| | | Geographic spec. | **0.657** | **0.640** | **0.630** | **0.648** | **0.641** | **0.645** | **0.636** | 3 |
| | With background strata | Random | **0.651** | **0.640** | **0.632** | **0.641** | **0.641** | **0.652** | **0.632** | 2 |
| | | Prop.-stratified | 0.603 | 0.594 | 0.592 | 0.597 | 0.589 | 0.596 | 0.594 | 7 |
| | | Equal-stratified | 0.597 | 0.588 | 0.586 | 0.591 | 0.583 | 0.589 | 0.590 | 8 |
| | | Density | **0.650** | **0.638** | **0.631** | **0.640** | **0.639** | **0.651** | **0.629** | 3 |
| | | Target | 0.638 | 0.627 | 0.621 | 0.627 | 0.629 | 0.639 | 0.617 | 5 |
| | | Density spec. | 0.645 | 0.629 | 0.628 | 0.635 | 0.625 | 0.637 | 0.620 | 4 |
| | | Target spec. | 0.627 | 0.612 | 0.610 | 0.616 | 0.608 | 0.624 | 0.600 | 6 |
| | | Geographic spec. | **0.656** | **0.642** | **0.637** | **0.646** | **0.642** | **0.654** | **0.632** | 1 |

**Table S2.** SDM predictive performance (AUC metric, evaluated on the independent presence/absence dataset) of ensemble models assessed across 500 plant species under eight pseudo-absences sampling strategies (average among pseudo-absence replicates) for the ensemble and single models of five algorithms (glm, gam, gbm, rdf and max). The three best evaluations are highlighted in bold and the best pseudo-absences sampling strategies are ranked (Rank). glm = generalized linear models, gam = generalized additive model, gbm = gradient boosting machines, rdf = Random Forest, max = maximum entropy.

| Pseudo-absence selection strategy | | | Ensemble evaluation (AUC) | Single model evaluations (AUC) | | | | | | Rank |
|---|---|---|---|---|---|---|---|---|---|---|
| | | | | mean | glm | gam | gbm | rdf | max | |
| Without adding spatial bias | Without background strata | Random | **0.916** | **0.909** | **0.903** | **0.908** | **0.905** | **0.921** | **0.911** | 1 |
| | | Prop.-stratified | 0.887 | 0.880 | 0.877 | 0.883 | 0.869 | 0.890 | 0.882 | 7 |
| | | Equal-stratified | 0.883 | 0.876 | 0.874 | 0.879 | 0.864 | 0.885 | 0.878 | 8 |
| | | Density | **0.914** | **0.907** | **0.901** | **0.905** | **0.903** | **0.918** | **0.909** | 2 |
| | | Target | 0.901 | 0.895 | 0.887 | 0.890 | 0.891 | 0.908 | 0.896 | 4 |
| | | Density spec. | 0.890 | 0.878 | 0.871 | 0.876 | 0.876 | 0.885 | 0.883 | 6 |
| | | Target spec. | 0.897 | 0.887 | 0.880 | 0.882 | 0.883 | 0.898 | 0.893 | 5 |
| | | Geographic spec. | **0.911** | **0.903** | **0.896** | **0.901** | **0.900** | **0.913** | **0.905** | 3 |
| | With background strata | Random | **0.912** | **0.905** | **0.899** | **0.903** | **0.900** | **0.918** | **0.906** | 1 |
| | | Prop.-stratified | 0.882 | 0.875 | 0.868 | 0.873 | 0.866 | 0.889 | 0.877 | 7 |
| | | Equal-stratified | 0.878 | 0.870 | 0.863 | 0.868 | 0.862 | 0.885 | 0.873 | 8 |
| | | Density | **0.911** | **0.905** | **0.898** | **0.902** | **0.900** | **0.917** | **0.905** | 2 |
| | | Target | 0.902 | 0.896 | 0.888 | 0.891 | 0.893 | 0.909 | 0.896 | 4 |
| | | Density spec. | 0.898 | 0.889 | 0.883 | 0.887 | 0.886 | 0.900 | 0.888 | 6 |
| | | Target spec. | 0.901 | 0.894 | 0.887 | 0.890 | 0.892 | 0.908 | 0.895 | 5 |
| | | Geographic spec. | **0.910** | **0.902** | **0.896** | **0.901** | **0.899** | **0.915** | **0.903** | 3 |
| With adding spatial bias | Without background strata | Random | **0.894** | **0.886** | **0.882** | **0.886** | **0.886** | **0.891** | **0.887** | 1 |
| | | Prop.-stratified | 0.867 | 0.860 | 0.861 | 0.864 | 0.854 | 0.857 | 0.865 | 6 |
| | | Equal-stratified | 0.864 | 0.857 | 0.859 | 0.861 | 0.849 | 0.852 | 0.862 | 7 |
| | | Density | **0.893** | **0.885** | **0.881** | **0.885** | **0.885** | **0.890** | **0.885** | 2 |
| | | Target | 0.883 | 0.876 | 0.872 | 0.876 | 0.876 | 0.881 | 0.875 | 4 |
| | | Density spec. | 0.875 | 0.859 | 0.841 | 0.867 | 0.866 | 0.855 | 0.865 | 5 |
| | | Target spec. | 0.836 | 0.824 | 0.811 | 0.809 | 0.835 | 0.825 | 0.839 | 8 |
| | | Geographic spec. | **0.891** | **0.880** | **0.871** | **0.882** | **0.882** | **0.883** | **0.882** | 3 |
| | With background strata | Random | **0.889** | **0.882** | **0.877** | **0.881** | **0.882** | **0.888** | **0.881** | 2 |
| | | Prop.-stratified | 0.863 | 0.857 | 0.855 | 0.858 | 0.852 | 0.857 | 0.861 | 7 |
| | | Equal-stratified | 0.859 | 0.853 | 0.852 | 0.855 | 0.848 | 0.853 | 0.858 | 8 |
| | | Density | **0.888** | **0.881** | **0.877** | **0.881** | **0.881** | **0.887** | **0.881** | 3 |
| | | Target | 0.882 | 0.875 | 0.871 | 0.874 | 0.875 | 0.881 | 0.873 | 5 |
| | | Density spec. | 0.886 | 0.877 | 0.875 | 0.878 | 0.874 | 0.881 | 0.874 | 4 |
| | | Target spec. | 0.875 | 0.866 | 0.865 | 0.868 | 0.864 | 0.871 | 0.863 | 6 |
| | | Geographic spec. | **0.891** | **0.883** | **0.880** | **0.883** | **0.883** | **0.889** | **0.881** | 1 |
